# Fluorescein-Based SynNotch Adaptors for Regulating Gene Expression Responses to Diverse Extracellular Cues

**DOI:** 10.1101/2024.06.12.598538

**Authors:** Jeremy C. Tran, Christopher J. Kuffner, Alexander M. Marzilli, Ryan Emily Miller, Zachary E. Silfen, Jeffrey B. McMahan, D. Christopher Sloas, Christopher S. Chen, John T. Ngo

## Abstract

We introduce an adaptor-based strategy for regulating fluorescein-binding synthetic Notch (SynNotch) receptors using ligands based on conjugates of fluorescein isomers and analogs. To develop a versatile system, we evaluated the surface expression and activities of multiple constructs containing distinct extracellular fluorescein-binding domains. Using an optimized receptor, we devised ways to regulate signaling via fluorescein-based chemical transformations, including an approach based on a bio-orthogonal chemical ligation and a spatially controllable strategy via the photo-patterned uncaging of an *o*-nitrobenzyl-caged fluorescein conjugate. We further demonstrate that fluorescein-conjugated extracellular matrix (ECM)-binding peptides can regulate SynNotch activity depending on the folding state of collagen-based ECM networks. Treatment with these conjugates enabled cells to distinguish between folded versus denatured collagen proteins and enact dose-dependent gene expression responses depending on the nature of the signaling adaptors presented. To demonstrate the utility of these tools, we applied them to control the myogenic conversion of fibroblasts into myocytes with spatial and temporal precision and in response to denatured collagen-I, a biomarker of multiple pathological states. Overall, we introduce an optimized fluorescein-binding SynNotch as a versatile tool for regulating transcriptional responses to extracellular ligands based on the widely used and clinically-approved fluorescein dye.

## INTRODUCTION

Multicellular processes, including development and tissue regeneration, require precise signaling coordination between cells and their surroundings. For example, Notch signaling permits cells to regulate gene expression according to the positional and contextual information they receive via direct cell-cell contacts^1,2^. Similarly, changes in interactions with extracellular matrix (ECM) via integrin receptors can trigger diverse cellular processes, ranging from cytoskeletal reorganization and adhesion to stem cell differentiation^3^. Inspired by these natural mechanisms, researchers have engineered ways to control and reprogram how cells interact with their microenvironments. For example, ‘bio-instructive materials’ have been developed to mimic natural ECM properties, and such materials have been used to direct the healing and repair of damaged tissues^4,5^. In recent years, tailored biomaterials have also been developed to control engineered cell activities, and these systems have been applied to modulate therapeutic T-cell responses *in situ* and to facilitate immune cell engineering *in vivo*^6–9^. Yet despite these advances, synthetic biology strategies for programming customized ECM-dependent signaling activities have remained relatively limited, especially in comparison to tools for sensing and programming responses to cell-cell interactions^10–14^.

Recognizing the engineering utility of cell-ECM signaling components, we aimed to create a versatile approach that could be used to define how cells sense and interpret scaffold-embedded signals within their surroundings. As a basis for this toolset, we exploited the modularity of the synthetic Notch (SynNotch) design, an engineered receptor framework that can link user-specified ligands to customized gene transcription responses^10,15^. Like natural Notch, SynNotch receptors contain intracellular domains (ICDs) based on transcriptional effector proteins. During signaling, these effectors are proteolytically cleaved from their transmembrane sequences to enable their nuclear translocation for the regulation of target genes. For both native and synthetic Notch, such cleavages require the tension-mediated unfolding of a juxtamembrane segment termed the Negative Regulator Region (NRR)^16,17^. In natural contexts, this unfolding is mediated via the tensile forces delivered to Notch receptors during their *trans*-endocytosis into ligand-expressing cells^16,18^. Soluble ligands cannot generate the required tensile energies and therefore often serve as competitive signaling inhibitors.

Surface and bead-immobilized ligands can also facilitate tension-mediated Notch and SynNotch activation, and by exploiting such ligands, researchers have devised ways to elicit desirable signaling responses from various cell types, *in vitro* and *in vivo*^19–23^. Furthermore, recent work has shown that ligands bound to endogenous and engineered material scaffolds can also facilitate productive signaling. For example, immobilization of the Delta-like 1 (DLL1) ligand has been applied to activate Notch in cells embedded within photopatterned fibrin-based hydrogels^24^, and a bone mineral-binding ligand has been used to selectively stimulate Notch in cells surrounding skeletal growth plates *in vivo*^25^. From a synthetic biology perspective, SynNotch receptors recognizing native ECM components have recently been developed using a collagen-II binding antibody fragment^26^, and strategies involving microcontact printed ligands have been exploited to define SynNotch activities with micro-scale spatial precision^27,28^.

Inspired by the growing repertoire of Notch and SynNotch activation strategies, we set out to devise an adaptor-based strategy that could be used to gain versatile and dynamic control over SynNotch signaling via a ‘universal’ ligand-receptor pair. We reasoned that with such components, one could bypass the need to individually construct and characterize new receptor extracellular domains---the engineering of which can be bespoke with challenges associated with converting IgGs into well-behaved single-chain sequences. Building upon the previous design of ‘universal’ chimeric antigen receptors (CARs)^29–33^, we describe a fluorescein-based strategy for regulating the activity of a fluorescein-binding SynNotch with inducible and dose-dependent control. To develop this approach, we first characterized SynNotch constructs fused with distinct fluorescein-binding domains, analyzing their surface levels and quantifying their activities against ligands based on fluorescein isomers and analogs. Using an optimized receptor, we defined conditions to maximize ligand-induced gene expression responses while minimizing background (ligand-independent) signaling activities. Roles for the presenilin isoforms (presenilin-1 and presenilin-2) were also investigated in the context of ligand-mediated and ligand-independent receptor activities, and cells expressing our optimized construct were used to define the kinetics of ligand-induced reporter mRNA formation and fluorescent protein expression.

To highlight the versatility of our system, we exploited fluorescein-based tools to regulate SynNotch signaling in biocompatible and spatially controllable ways. Using bio-orthogonal chemistry, we show that tetrazine-functionalized fluorescein can be transformed into a signaling active ligand via its selective ligation with immobilized *trans*-cyclooctene groups in the presence of live cells. To achieve spatial control over signaling, we employed photo-caged fluorescein as a light-conditional ligand, using photo-masked illumination to generate spatially defined ligand patterns in 2D. Additionally, we exploit fluorescein-conjugated ECM-binding peptides to direct ECM-dependent SynNotch activities in response to native and unfolded collagen-I proteins. Finally, to demonstrate the utility of these tools, we exploited them to regulate the myogenic conversion of multipotent fibroblasts into myoblasts with spatiotemporal control and molecular precision.

## RESULTS

### Fluorescein-binding SynNotch receptors are correctly processed and trafficked to the cell surface

To develop a sensitive and versatile system, we sought an efficiently expressed receptor capable of detecting ligands based on commonly-used fluorescein isomers and analogs. Synthetic receptors were generated by fusing fluorescein-binding domains to the extracellular region of the SynNotch scaffold (**Figure 1a**). Three previously characterized domains were tested in our designs, including two single-chain variable fragments (scFvs) based on existing anti-fluorescein antibodies—*α*FITC(E2)^34,35^ and 4M5.3^36,37^—and a third (non-immunoglobulin) domain based on the engineered lipocalin “FluA”^38–40^(**Figure 1b**). The resulting fusions are hereafter referred to as *α*FITC(E2)-SynNotch, 4M5.3-SynNotch, and FluA-SynNotch, respectively. Beyond their distinct binding domains, each construct contained otherwise identical components, including a CD8*α* signal peptide, an extracellular myc-tag (for surface detection), and a core domain based on the Notch1 NRR and its transmembrane helix. We used receptors containing an intracellular domain (ICD) based on Gal4-VP64 to quantify ligand-induced signals in cells containing Gal4-dependent reporter genes.

**Figure 1.**
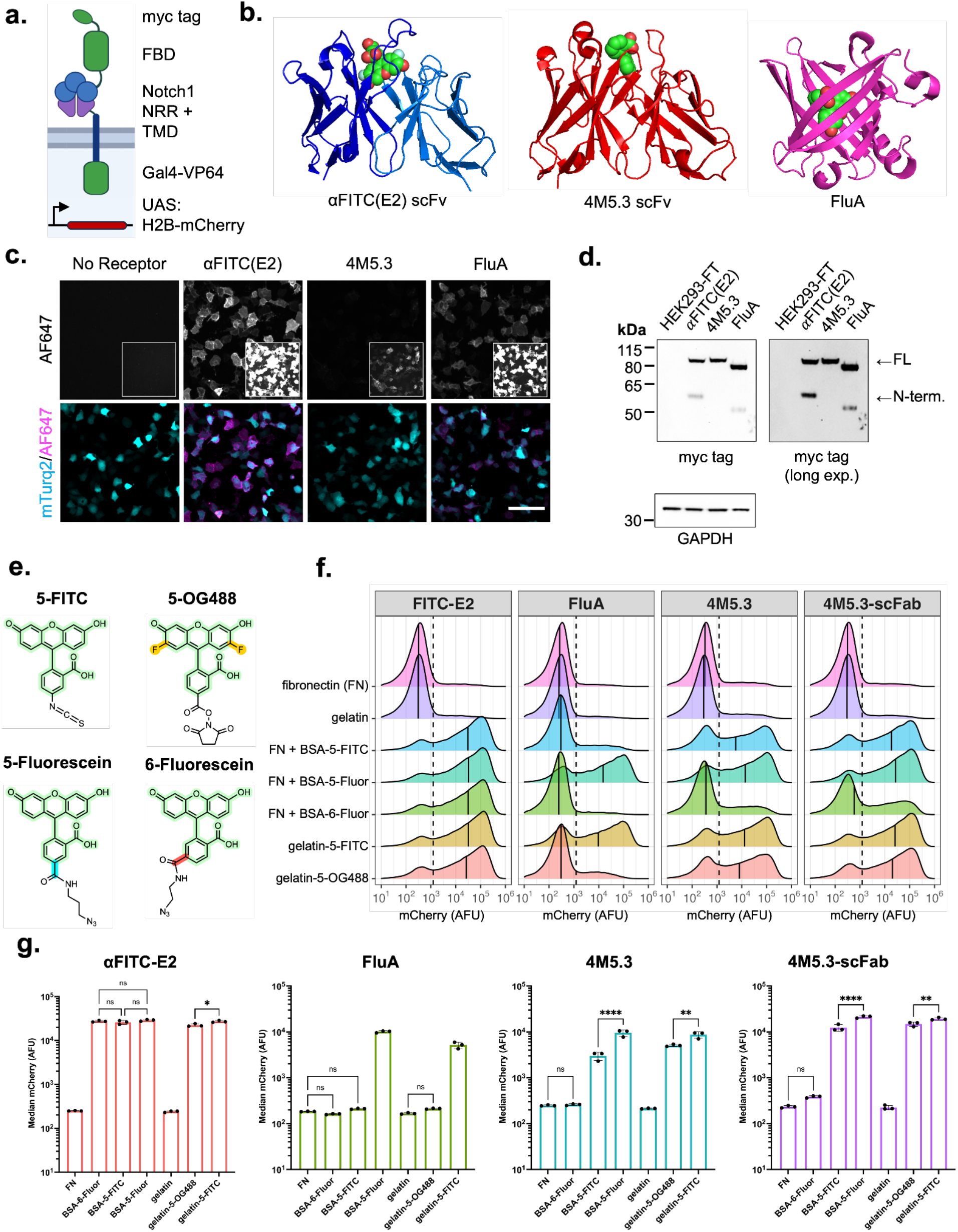
Characterization of fluorescein-binding SynNotch receptors. (**a**) Schematic depicting the design SynNotch receptors containing the various tested fluorescein binding domains (FBDs). (**b**) Dye-bound structures of the various FBDs tested in our designs, including *α*FITC(E2) (PDB: 2A9N, a bound to the difluorinated fluorescein analog, OregonGreen488), 4M5.3 (PDB: 1X9Q), and FluA (PDB: 1N0S). (**c**) Detection of surface-localized SynNotch receptors expressed on transfected HEK293-FT cells. Cells were co-transfected with DNAs encoding the indicated receptors in combination with a mTurq2-encoding plasmid as a co-transfection marker. Cells were immunostained the next day under live-cell conditions using an anti-myc-AlexaFluor647 (anti-myc-AF647) antibody conjugate. Insets represent over-exposed AF647 emissions. Scale bar = 100 μm. (**d**) Immunoblot detection of myc-tagged receptors. Whole-cell lysates from transfected HEK293-FT cells were analyzed. Bands corresponding to full-length (FL, higher mass) and furin-cleaved N-terminal (N-term., lower mass) receptor fragments are indicated with arrows. In both (c) and (d), HEK293-FT cells transfected with an mTurq2-encoding plasmid were analyzed as controls to confirm the specificity of the utilized anti-myc antibody probes. (**e**) Structures of the fluorescein isomers and analogs used as test ligands against the various receptor designs. (**f**) Flow cytometry traces of reporter expression levels in transfected HEK293-FT (UAS:H2B-mCherry) reporter cells expressing the indicated receptors. The transfected cells were grown overnight in microwells containing the indicated ligands and adhesion proteins. Ligand stimulation proceeded overnight prior to flow cytometry analyses; wells were pre-coated the indicated ligands prior to adding cells (see Methods). Traces represent normalized densities of three independent transfections (n=3, > 5,000 cells analyzed per replicate) gated for receptor expression by the mTurq2 co-transfection marker. The dashed black line indicates the fluorescence threshold used to define H2B-mCherry+ cells, as determined via analysis of non-transfected HEK293-FT cells; solid black lines indicate median mCherry intensities for each depicted population. (**g**) Median mCherry intensities from mTurq2+ population from (f); Values are analyzed with two-way ANOVA (ligand and receptor). All unlabelled comparisons involving fibronectin and gelatin controls versus co-plated ligands have P < 0.0001; labeled NS, P>0.05, *P < 0.05, **P<0.01, ***P<0.001, ****P<0.0001, otherwise. Fluor; fluorescein, OG488; OregonGreen488.

Since efficient surface expression is required for sensitive extracellular ligand detection, we first evaluated the presentation of our constructs using transfected HEK293-FT cells. Immunolabeling with a fluorescent anti-myc antibody revealed varying cell surface levels for our designs: *α*FITC(E2)-SynNotch was the most efficiently presented, followed by FluA-SynNotch, which was moderately reduced by comparison. 4M5.3-SynNotch had the lowest surface levels among the evaluated receptor sequences (**Figure 1c, Supplementary Figure 1a**). Furin is a Golgi-localized enzyme that processes the NRR into a heterodimeric complex before the receptors are presented at the plasma membrane^41,42^, and blotting of cell lysates showed that full-length receptors were translated to similar levels but were processed by furin to varying degrees (**Figure 1d**). The lack of a furin-cleaved band for 4M5.3-SynNotch suggested its retention within the endoplasmic reticulum (ER), which we hypothesized to be due to misfolding of the 4M5.3 scFv. Correspondingly, converting the scFv into the more stable single-chain Fab (scFab) format^43^ resulted in 4M5.3-scFab-SynNotch, which was efficiently cleaved by furin and trafficked to the surface in multiple transfected cell lines (including HEK293-FT, U2OS, and CHO-K1 cells, **Supplemental Figure 1b-c**). Thus, *α*FITC(E2)-SynNotch and FluA-SynNotch are efficiently trafficked constructs, and our data show that the trafficking inefficiency of 4M5.3-SynNotch can be overcome via substitution of the scFv with a 4M5.3-based scFab. Similar manipulations may help improve the display of other scFv-based antigen-binding receptor systems.

### *α*FITC(E2)-SynNotch can sensitively detect ligands based on immobilized fluorescein

Next, we evaluated the signaling activities of our receptors using a model ligand based on fluorescein-5-isothiocyanate (FITC)-conjugated bovine serum albumin (BSA-5-FITC). Reporter cells containing a Gal4-dependent gene (HEK293-FT:UAS-H2B-mCherry) were transiently transfected with receptor-encoding plasmids and grown overnight in BSA-5-FITC-coated microwells. Flow cytometry quantification of H2B-mCherry levels confirmed the activation of our receptors by the immobilized BSA-5-FITC, with controls showing that such signals were fluorescein-dependent (**Figure 1e-g**). Dose-dependent analyses with BSA-5-FITC showed that *α*FITC(E2)-SynNotch produced the most sensitive responses, followed by 4M5.3-scFab-SynNotch, 4M5.3-SynNotch, and FluA-SynNotch, respectively (**Supplementary Figure 2**). Signaling levels in 4M5.3-scFab-SynNotch cells were higher than those expressing the scFv-containing 4M5.3-SynNotch, likely due to the increased surface levels of the scFab-containing design. To investigate the kinetics of signal activation, we directly probed time-dependent reporter mRNA formation using the hybridization chain reaction (HCR)^44^. Nuclear intensities corresponding to nascent *mCherry* transcripts were detected within 1-2 hours following ligand-induced ICD release (**Supplementary Figure 3**). These results show that *α*FITC(E2)-SynNotch activation and its downstream reporter gene expression proceed along timescales consistent with those reported for Notch nuclear complex assembly and gene transcription^45–47^. Together, these results confirm the fluorescein-dependent activities of our receptor designs and provide insight into the timescales of synthetic signaling-induced transcript formation.

### *α*FITC(E2)-SynNotch facilitates a versatile detection of fluorescein isomers and analogs

Fluorescein bioconjugates (including antibodies, etc.) can contain chemically distinct dye isomers and derivatives, some of which may not interact with the ligand binding domains used in our designs. Thus, to identify a highly-versatile receptor, we next tested our receptors against commonly used fluorescein isomers and analogs. We first analyzed responses to ligands based on 5-fluorescein versus 6-fluorescein—two widely used fluorescein isomers that differ in the positioning of where linker handles are attached to the carboxyphenyl ring (**Figure 1e**). Isomerically pure dyes were used to generate BSA-5-fluorescein and BSA-6-fluorescein conjugates (see **Methods**), and responses to these ligands were quantified as described above. Following overnight stimulation, measurement of H2B-mCherry levels revealed striking differences in the isomer binding preferences of our designs (**Figure 1f-g**). Most notably, FluA-SynNotch was activated by BSA-5-fluorescein but not the BSA-6-fluorescein, revealing a strict binding selectivity for the 5-isomer by the lipocalin-derived domain. Cells expressing 4M5.3-SynNotch and 4M5.3-scFab-SynNotch were heavily biased toward 5-fluorescein-based ligands, though low activity levels were detected in cells treated with BSA-6-fluorescein (**Figure 1f**).

In stark contrast to FluA-SynNotch cells, cells expressing *α*FITC(E2)-SynNotch produced potent responses to BSA-5-fluorescein and BSA-6-fluorescein, as well as a ligand based on the difluorinated fluorescein analog OregonGreen488^48^ (OG488, gelatin-OG488; **Figure 1e-g**). Together, our analyses revealed several advantageous properties for *α*FITC(E2)-SynNotch, including its efficient surface display and sensitive and unbiased recognition of fluorescein isomers and OG488. Given these advantages, we proceeded with *α*FITC(E2)-SynNotch, further evaluating its sensitivity to OG488 using cell-bead and cell-cell-based *trans*-activation assays. Here, “sender” cells expressing a biotinamide-binding ligand protein (anti-bio-SNAP-TMD-DLL1) were cocultured with *α*FITC(E2)-SynNotch “receivers,” and treatment with biotin-based bridging compounds was used to template the ligand-receptor interactions *in trans*^49^ (**Supplementary Figure 4**). Using this approach, we evaluated the sensitivity of bridges based on biotin-FITC versus biotin-OG488 conjugates, finding a more potent activity for biotin-OG488, which triggered 192-fold greater H2B-mCherry levels compared to biotin-FITC in response to 20 pM treatment concentrations (**Supplementary Figure 4f**). This result suggests OG488 could potentially be a more potent adaptor ligand for “universal” CAR designs involving the *α*FITC(E2) domain. Of note, treatment of cocultures with a short-linked fluorescein conjugate (biotin-ethylenediamine-fluorescein) did not lead to detectable reporter activities, suggesting that sufficient spacing between biotin and dye handles is needed for efficient templating of ligand-receptor complex between cells (**Supplementary Figure 4g**). In addition to cell-mediated *trans-*activation, treatment with fluorescein-and OG488-decorated microbeads also resulted in potent signaling activities (**Supplementary Figure 5**). Overall, these results highlight the versatility and sensitivity of *α*FITC(E2)-SynNotch to fluorescein isomers and analogs.

### Expression levels influence ligand-dependent and -independent receptor activities

Receptor overexpression can lead to ligand-independent signaling, limiting the dynamic range of *bona fide* signaling responses^50^. Thus, to facilitate the reliable adaptation of our tools, we optimized expression conditions needed to achieve low background and high receptor inducibility in using transfected and transduced reporter cells. First, HEK293-FT reporter cells were transfected with varying amounts of receptor-encoding plasmid (“pcDNA3-*α*FITC(E2)-LaG17-SynNotch-Gal4VP64”), and we evaluated H2B-mCherry following overnight incubation in BSA-5-FITC-coated versus uncoated microwells (**Supplementary Figure 6**). By comparing reporter levels between ligand-treated versus untreated cells, we identified 25 ng of the pcDNA3-based plasmid as an optimal amount for transfecting 150,000 HEK293-FT reporter cells (see **Methods**). Under these conditions, ligand stimulation led to a 191-fold increase in H2B-mCherry expression compared to untreated (receptor-expressing) reporters. Increased plasmid levels led to higher ligand-independent reporter levels, whereas lower amounts reduced ligand-induced reporter yields. In contrast to transient transfection, receptor expression by lentiviral transduction resulted in potently inducible cells without discernible increases in background activities over a 20-fold viral supernatant dose range (**Supplementary Figure 7**).

### Ligand-mediated *α*FITC(E2)-SynNotch signaling mediated primarily by presenilin-1 gamma-secretase (PS1-GS) in HEK293-FT cellss

Work by others has implicated gamma-secretase (GS) cleavage as a key mediator of the ‘leaky’ (ligand-independent) activity of cells expressing SynNotch receptors in excess^50^. GS is a multi-subunit enzyme composed of APH1, PEN-2, and nicastrin subunits in combination with a catalytic subunit based on one of two presenilin (PS) isoforms--PS1 or PS2, encoded by *PSEN1* and *PSEN2*, respectively^51^. To dissect the contributions of PS1 and PS2 in SynNotch processing, we used Cas9-editing to individually and doubly knockout (KO) PS expression in our HEK293-FT reporter cells (generating PS1-KO, PS2-KO, and PS1/PS2-double KO (DKO) lines) (**Supplementary Figure 8a**). Signaling analyses in the cells showed that ligand-induced activities were substantially diminished in PS1-lacking cells, whereas those of PS2-KO and non-KO cells were comparable (**Supplementary Figure 8b-e**). This result suggests a more prominent role for PS1-GS proteolysis in the ligand-induced activation of *α*FITC(E2)-SynNotch in HEK293-FT cells. In contrast, ligand-independent activities proceeded to comparable levels across individual presenilin KO and non-KO cells, being diminished only in PS1/PS2 DKO cells (**Supplementary Figure 8d**,**f**). Thus, ligand-induced SynNotch activities arise primarily due to PS1-mediated cleavage in HEK293-FT cells, whereas ligand-independent signals can derive from either PS1-GS, PS2-GS, or both enzymes to comparable degrees.

### Inducible signaling via bioorthogonal bond-formation between tetrazine and trans-cyclooctene handles

Having characterized *α*FITC(E2)-SynNotch, we next exploited fluorescein-based ligands to devise new methods for inducible receptor activation and gene expression control. Given that immobilized ligands can activate signaling, we predicted that soluble fluorescein units could be converted to signaling-competent ligands using cell-compatible bioorthogonal ligation reactions.

To test this possibility, we exploited bioorthogonal chemical ligation between tetrazine (Tz) and *trans*-cyclooctene (TCO) handles to immobilize a Tz-functionalized fluorescein dye to microwells containing an immobilized TCO-BSA conjugate (**Figure 2a-b**). Tz and TCO groups can react via an ultrafast inverse electron–demand Diels–Alder ligation reaction compatible with live cells and *in vivo* conditions^52,53^. Consistent with our expectations, Tz-5-fluorescein treatment triggered dose-dependent reporter activities upon its addition to 2D cell cultures grown in TCO-BSA-coated microwells. Measurement signaling levels showed that 2 nM treatment doses triggered a 487x-fold change in H2B-mCherry levels compared to Tz-5-fluorescein-untreated cells grown in TCO-BSA wells (**Figure 2c-f**). In contrast, cells grown without TCO-BSA were refractory to Tz-5-fluorescein treatment. In live cell time-lapse imaging, ligand treatment led to synchronous and uniform cell activity across TCO-BSA-containing wells (**Supplemental Movies 1-2**). These results show that bioorthogonal chemistry can be exploited to convert soluble fluorescein dyes into immobilized (and thus signaling-active) ligand handles.

**Figure 2.**
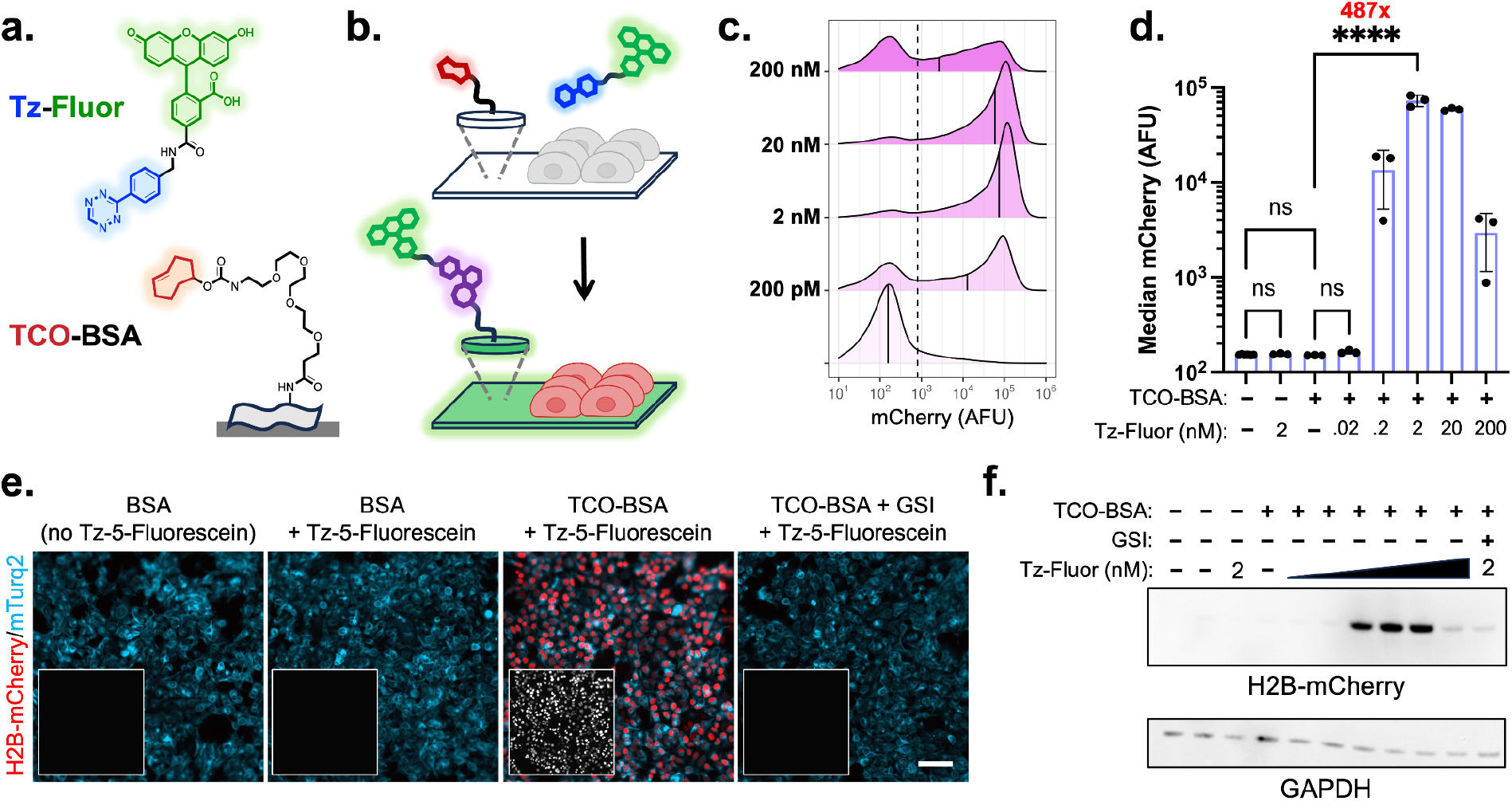
Receptor activation via a bioorthogonal Tz-TCO ligation reaction. (**a**) Chemical structures of Tetrazine-5-Fluorescein (Tz-5-fluorescein) and the *trans*-cyclooctene (TCO) reactive handle depicted as conjugated to BSA (TCO-BSA). (**b**) Schematic depiction of the bio-orthogonal ligation-mediated immobilization of Tz-5-fluorescein upon its reaction to surface-adsorbed TCO-BSA. Generation of the immobilized fluorescein ligand results in productive *α*FITC(E2)-SynNotch activation and H2B-mCherry reporter expression. (**c**) Dose-dependent analysis of Tz-5-fluorescein-mediated H2B-mCherry expression from *α*FITC(E2)-SynNotch cells grown in TCO-BSA coated microwells. Traces represent normalized densities of three independent treatments at varying doses are shown. The dashed black line indicates the H2B-mCherry+ threshold set based on analysis of non-transfected HEK293-FT cells; solid black lines indicate median mCherry intensities for each condition. (**d**) Median mCherry emission intensities from untreated and Tz-5-fluorescein-treated cells grown with or with immobilized TCO-BSA. Displayed values were analyzed with two-way ANOVA (ligand and drug concentration). Unlabelled comparisons between Tz-Fluor concentrations on TCO-BSA ligand have P < 0.05; labeled NS, P>0.05, *P < 0.05, **P<0.01, ***P<0.001, ****P<0.0001 otherwise. (**e**) Fluorescence images of clonal HEK293-FT (UAS:H2B-mCherry) reporter cells expressing *α*FITC(E2)-SynNotch-mTurq2, as treated under the indicated conditions. Scale bar = 100 μm. (**f**) Immunoblot detection of signaling-induced H2B-mCherry levels. Tz-fluorescein concentrations were tested between 2 pM and 200 nM, with variation between adjacent lanes by a factor of 10-fold. For both (f) and (e), treatment with gamma secretase inhibitor (GSI, DAPT at 10 μM) led to diminished Tz-5-fluorescein-induced reporter H2B-mCherry levels.

### Light-inducible signaling using a photocaged fluorescein ligand

We next asked whether fluorescein-based ligands could be used to define spatial gene expression patterns in our engineered cells. To test this possibility, we implemented a photocaged (PC)-fluorescein as a light-conditional ligand, anticipating that photolytic uncaging of the compound could be used to control receptor binding and, as a result, define gene expression patterns in 2D (**Figure 3a-c**). Using BSA conjugated with a bis-5-carboxymethoxy-2-nitrobenzyl (bis-CMNB) modified 5-fluorescein (BSA-PC-5-fluorescein), we compared signaling responses to the caged (light-protected) and photo-uncaged versions of the compound. Flow cytometry, imaging, and western blotting analyses confirmed our expectations, showing that the uncaged ligand, but not its caged precursor, could activate *α*FITC(E2)-SynNotch signaling and induce H2B-mCherry expression in our HEK293-FT reporter line (**Supplemental Figure 9**).

**Figure 3.**
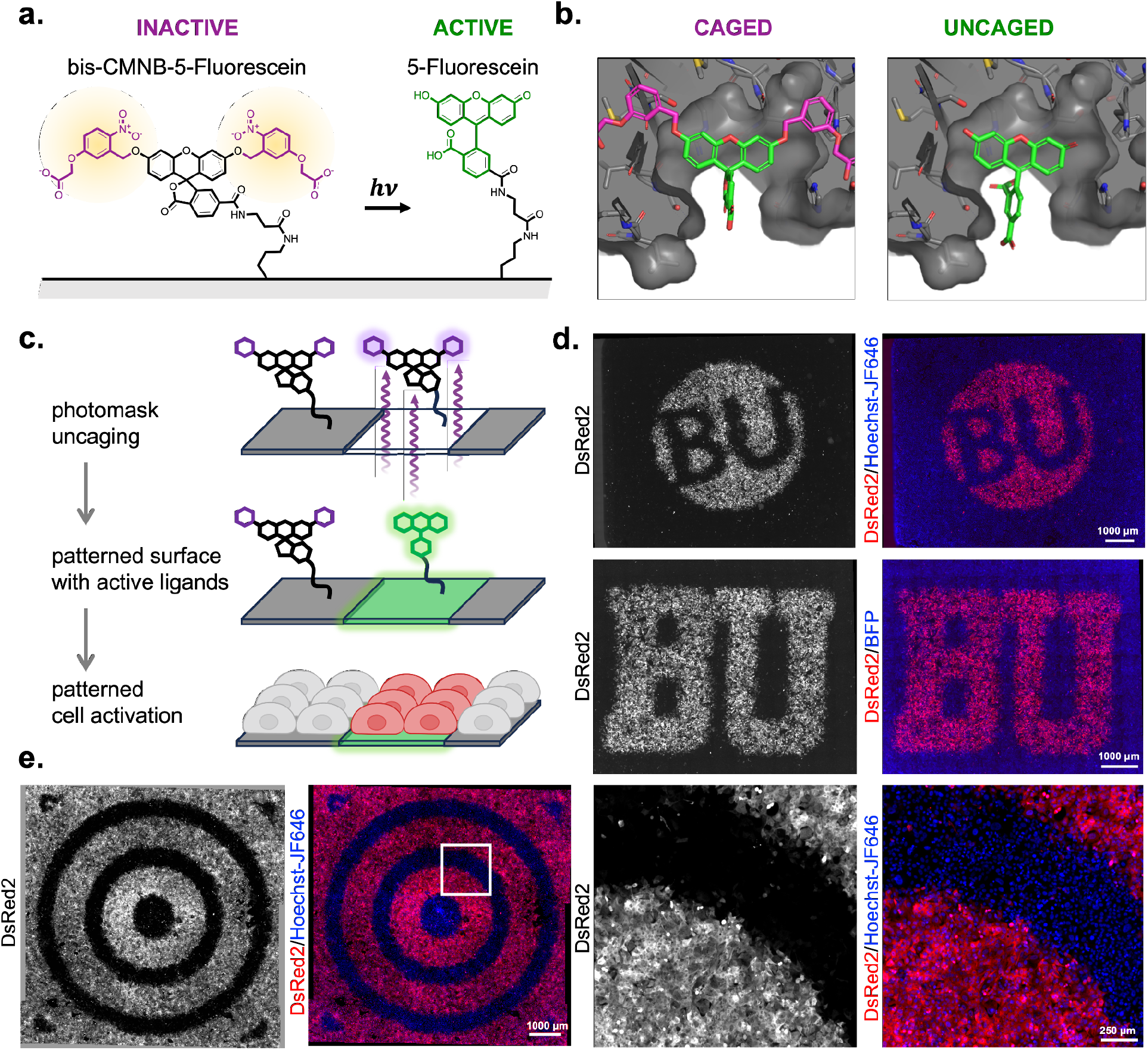
Gene expression patterning via photolytic conversion of a photo-caged fluorescein ligand. (**a**) Structure of photolabile bis-5-carboxymethoxy-2-nitrobenzyl (bis-CMNB) modified 5-fluorescein (PC-5-fluorescein) and UV-light induced photo-uncaged product 5-fluorescein. (**b**) Models of *α*FITC(E2) scFv binding (left) PC-5-carboxy-fluorescein; photolabile caging groups (purple) are hypothesized to prevent ligand binding. (right) 5-carboxy-fluorescein. Models were built by superimposing fluorescein derivative on OG488 ligand in PDB: 2A9N (**c**) Schematic of photo-uncaging plate adsorbed PC-5-Fluorescein leading to *α*FITC(E2)-SynNotch activation in defined areas. (**d-e**) Laser-printed transparency slides were used as photomasks to generate patterns of uncaged BSA-PC-5-fluorescein ligands. *α*FITC(E2)-SynNotch-T2A-BFP U2OS cells with DsRed2 reporter were seeded and imaged 24-36 h later.

Equipped with a light-conditional ligand, we next asked whether we could exploit BSA-PC-5-fluorescein to generate spatially defined gene expression patterns. Here, we devised simple photomasks based on laser-printed transparency slides, exploiting them to create uncaged PC-5-fluorescein-BSA patterns via transillumination with a handheld UV lamp (see **Methods**). U2OS cells expressing *α*FITC(E2)-SynNotch and containing a UAS:DsRed2 reporter construct were grown on the patterned surfaces, and fluorescence microscopy was used to visualize reporter expression patterns across 2D monolayers 24 - 36 hours after cell seeding (**Figure 3d-e, Supplementary Figure 10**). These analyses confirmed the spatial specificity of our approach, showing that reporter activities were confined to culture areas containing photo-uncaged fluorescein ligands.

We also examined ligand photoactivation in the presence of live cells using time-lapse imaging. Here, reporter activity was recorded following pulse photo-uncaging through a DAPI excitation filter (361 - 389 nm) (**Supplementary Movie 3**). Subsequent monitoring of reporter activity confirmed the confinement of signaling responses to illuminated well areas. To compare the time scales of SynNotch-mediated versus chemically-induced gene expression, we conducted time-dependent analyses using U2OS cells containing a TRE3G-mCherry reporter gene. Ligand photo-uncaging was used to activate a receptor containing a TetR-VP48 ICD, and time-lapse imaging was used to analyze reporter expression rates with direct comparison to doxycycline-mediated mCherry expression (via a co-expressed TetON-3G *trans*-activator protein). In both cases, mCherry expression was detected within 4-6 hours following light exposure (1 sec) or doxycycline treatment (200 ng/mL) (**Supplementary Figure 11**). These data confirm the live cell compatibility of ligand photo-uncaging, showing that light-mediated signal activation proceeds with comparable kinetics to doxycycline-induced gene expression. Overall, our results validate BSA-PC-5-fluorescein as a light-conditional ligand, showing that the caged molecule can be uncaged to direct spatially-defined gene expression in 2D cultures of SynNotch-expressing cells.

### Signal activation via ECM-binding bifunctional bridges

Next, we asked whether fluorescein-based agents could direct synthetic signaling responses to ECM substituent proteins. We reasoned that dye-conjugated ECM-binding proteins could be transformed into signaling-active ligands upon immobilization to ECM protein networks. To test this possibility, we generated an OG488-conjugated version of CNA35, a 35 kDa collagen-binding domain derived from the collagen adhesion protein of *Staphylococcus aureus*^54,55^ (**Figure 4a**). In cells grown on collagen-I substrates, OG488-CNA35 treatment induced dose-dependent *α*FITC(E2)-SynNotch activation in a manner that was selective for native collagen over gelatin (denatured collagen) and fibronectin (**Figure 4b-c**), consistent with CNA35’s well-characterized binding selectivity for folded collagen triple helices^56^.

**Figure 4.**
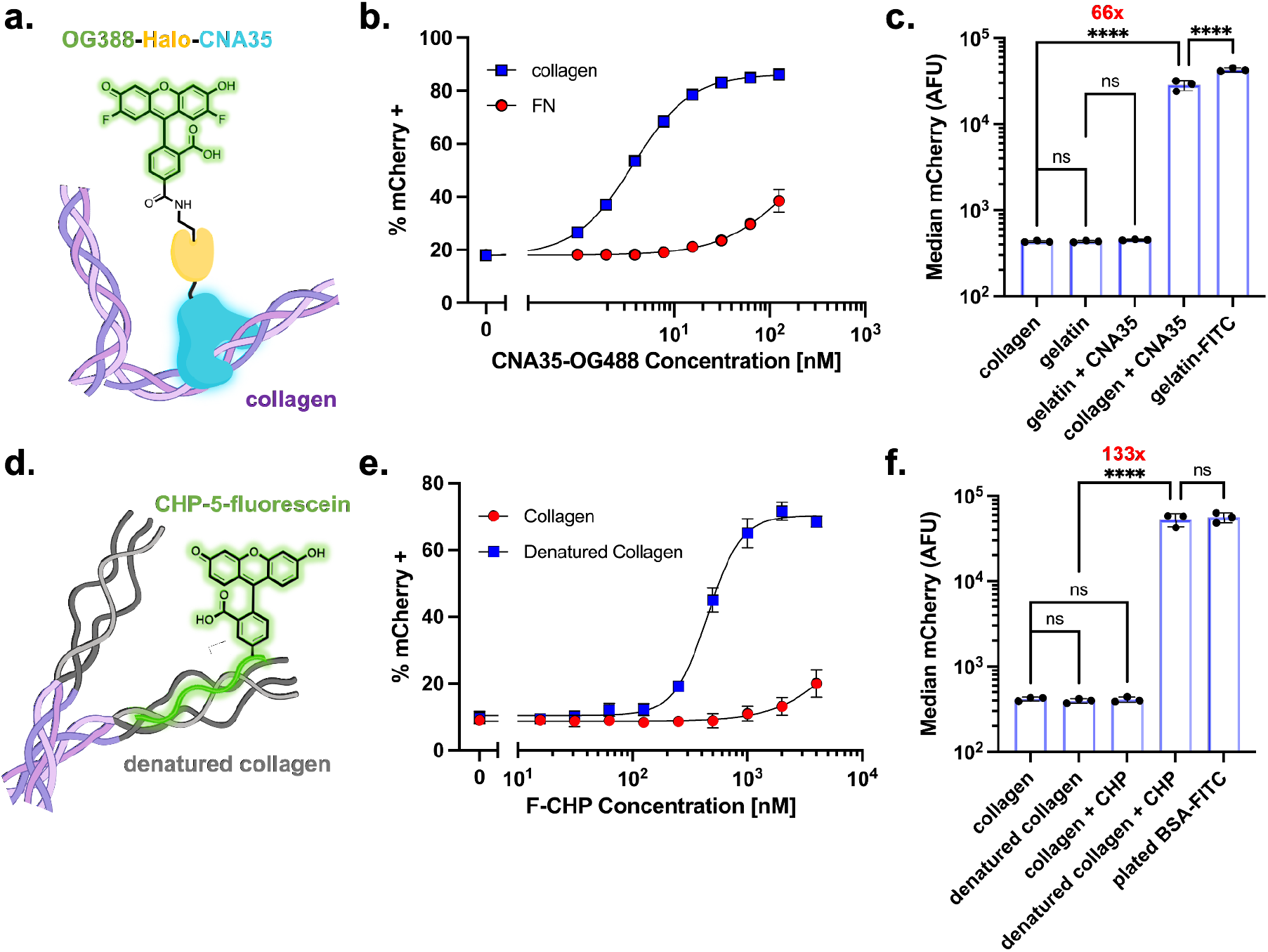
Receptor activation via dye-labeled collagen-binding proteins. **(a)** Schematic of OG488-CNA35 (CNA35 in blue, HaloTag in yellow, OG488 in green) bound to natively folded collagen-I (purple). **(b)** OG488-CNA35 activates *α*FITC(E2)-SynNotch specifically on collagen-I coated wells at low nanomolar concentration. HEK293-FT H2B-mCherry reporter cells were transiently transfected with plasmids encoding *α*FITC(E2)-SynNotch and mTurq2 as a cotransfection marker. Percent mCherry positive cells of the mTurq2+ population analyzed by flow cytometry from three independent transfections (n=3, > 5,000 cells per replicate) displayed. **(c)** Median mCherry fluorescence intensities showing clonal *α*FITC(E2)-SynNotch-mTurq2 cells specifically activated by 3 nM OG488-CNA35 bound to triple-helical collagen-I. Analyzed by one-way ANOVA, labeled NS: P > 0.05, ****: P <0.0001. **(d)** Schematic of CHP-5-fluorescein bound to denatured collagen (gray). **(e)** CHP-5-fluorescein activates on heat-denatured collagen-I. HEK293-FT H2B-mCherry reporter cells were transiently transfected with *α*FITC(E2)-SynNotch and mTurq2 cotransfection marker. Percent mCherry positive cells of the mTurq2+ population analyzed by flow cytometry from three independent transfections (n=3, >5,000 cells per replicate) displayed. **(f)** Median mCherry fluorescence intensities showing clonal *α*FITC(E2)-SynNotch-mTurq2 cells specifically activated by 2 μM CHP-5-fluorescein bound to heat-denatured collagen-I. Analyzed by one-way ANOVA, labeled NS: P > 0.05, ****: P <0.0001.

To complement the native-collagen sensing method above, we also designed a strategy to enable cells to detect damaged collagens in their microenvironments and activate synthetic gene expression as a response. Here, a fluorescein-conjugated version of the Collagen Hybridizing Peptide (CHP-5-fluorescein) was used as a bridging agent to immobilize receptor-binding fluorescein ligands to denatured collagen proteins. CHP is a short glycine-proline-hydroxyproline repeat that can hybridize with unfolded collagen *α*-chains^57^ and previous work has shown that CHP conjugates can be used to trace denatured collagens *in vivo*^58–60^ and to localize therapeutic molecules to diseased tissues^61,62^ (**Figure 4d**). Growth of *α*FITC(E2)-SynNotch cells on CHP-5-fluorescein-treated collagen substrates triggered dose-dependent responses to denatured collagen-I, with cells on native collagen producing only marginal responses, limited to wells treated with mid-micromolar-range peptide-conjugate concentrations (**Figure 4e-f**). Together, these results show that ligand-conjugated ECM-binding components can be exploited as ECM-receptor adaptors to enable cells to detect the composition and folding state of the ECM, directing them to activate synthetic gene transcription activities in response.

### Fluorescein-based ligands can facilitate the myogenic conversion of multipotent C3H10T1/2 cells

Finally, to demonstrate the utility of our tools, we implemented them to control the myogenic conversion of embryonic mesenchymal C3H10T1/2 cells containing a SynNotch-inducible p65-MyoD fusion gene (**Figure 5a-b**). MyoD is a master myogenic regulator, and p65-MyoD expression in C3H10T1/2 cells facilitates their efficient conversion into multinucleated syncytial myoblasts^27,49,63^. As a first test, we examined whether Tz-fluorescein treatment could trigger the myogenic conversion of C3H10T1/2 grown on TCO-BSA-coated substrates. Indeed, treatment with nanomolar Tz-fluorescein concentrations triggered DsRed2-expression and myoblast formation in the engineered fibroblasts in a TCO-BSA-dependent manner (**Figure 5c**).

**Figure 5.**
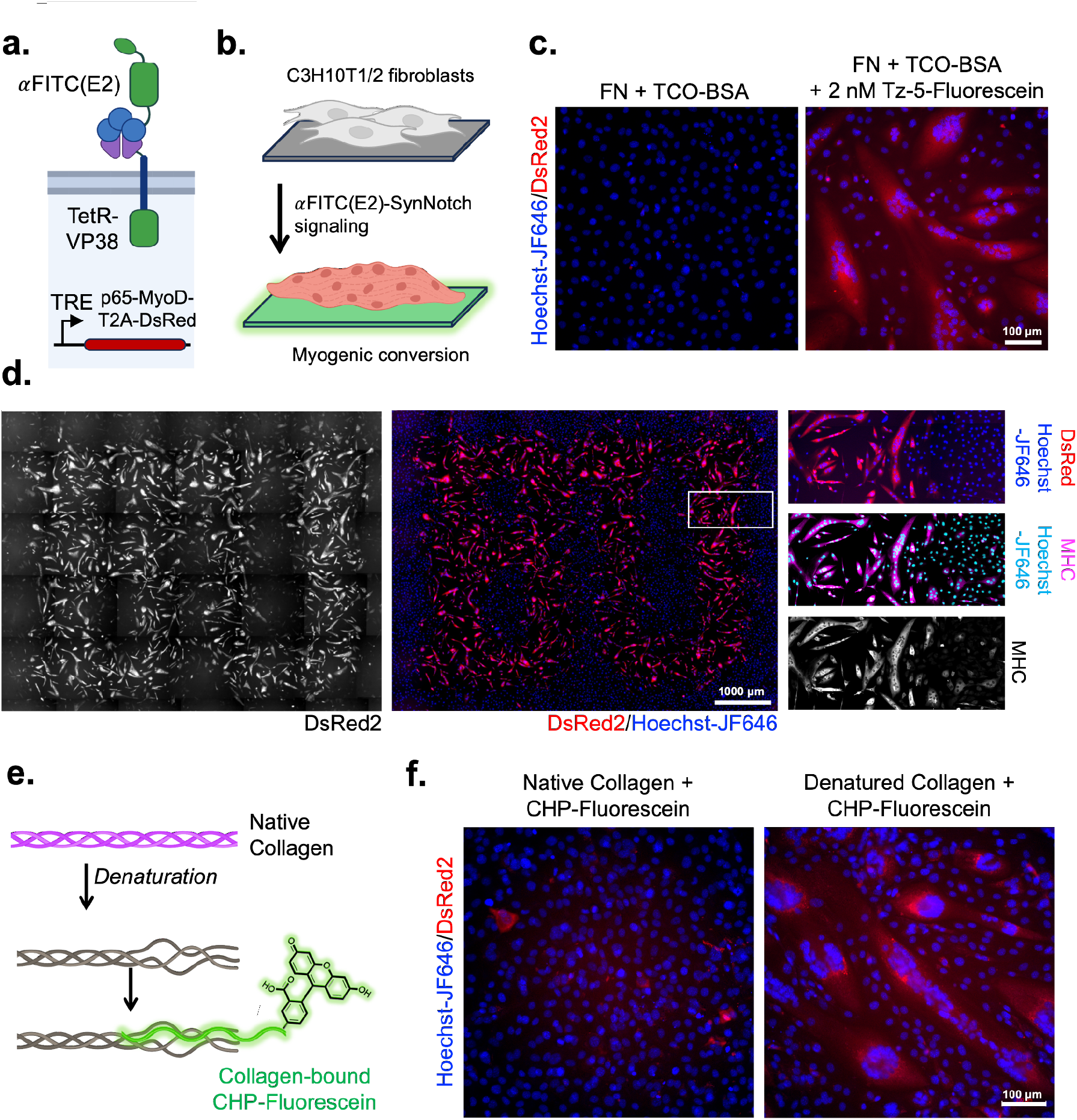
Conditional myogenic conversion by bioorthogonal ligation, photo-uncaging, and denatured collagen using fluorescein adapter-induced *α*FITC(E2)-SynNotch signaling. **(a)** Schematic depicting *α*FITC(E2)-SynNotch containing a TetR-VP48 ICD. The receptor is expressed in C3H10T1/2 fibroblasts containing a TRE-regulated target gene encoding p65-MyoD-T2A-DsRed2. **(b)** Depiction of the signaling-mediated conversion of C3H10T1/2 fibroblasts in response to fluorescein-induced *α*FITC(E2)-SynNotch activation. Expression of p65-MyoD-T2A-DsRed2 results in the formation of red fluorescent syncytia exhibiting myogenic phenotypes. **(c)** C3H10T1/2 fibroblasts grown in TCO-BSA coated wells were left untreated (left) or treated with 2 nM Tz-5-Fluorescein (right). Cells were imaged 48 hours following treatment. **(d)** C3H10T1/2 fibroblasts were grown in microwells containing immobilized and photo-patterned BSA-PC-5-fluorescein. Cells were imaged after 72 hours of growth on the patterned surfaces. Expression of the myogenic marker protein myosin heavy chains (MHC) was confirmed via immunofluorescence staining. **(e)** Schematic depicting the denaturation of collagen triple helices followed by the binding of CHP-fluorescein to immobilized collagen strands. **(f)** Microwells containing native (left) or denatured (right) collagen-I proteins were treated with CHP-fluorescein (2 μM) prior to the addition of C3H10T1/2 cells. Cells were imaged after 48 hours of growth.

Inspired by the highly organized alignment of skeletal myoblasts in natural muscle tissues, we tested the utility of photopatterned BSA-PC-5-fluorescein surfaces in directing myogenic conversion with spatial control. Following three days of growth in photopatterned microwells, cells in photo-activated well areas exhibited phenotypic markers of receptor activation and myogenesis, including DsRed2 fluorescence, multinucleation, and expression of the myogenic marker protein myosin heavy chain (MHC; **Figure 5d**). In contrast, cells residing in light-protected well regions remained in fibroblast-like states, as evident by their mononuclear morphologies and lack of anti-MHC immunoreactivity. Similar treatment of C3H10T1/2 cells containing a TRE:DsRed2 reporter gene (in place of TRE:p65-MyoD-T2A-DsRed2) resulted in confined DsRed2 expression patterns without discernible syncytia formation, verifying that the observed conversion of TRE:p65-MyoD-T2A-DsRed2 cells was due to signaling-mediated p65-MyoD expression (**Supplementary Figure 12**).

Finally, we asked whether treatment with CHP-5-fluorescein could be used to selectively direct the myogenic conversion of cells grown on denatured but not native collagen-I-based substrates. Consistent with our fluorescent reporter data, the growth of cells on denatured collagen in the presence of CHP-5-fluorescein led to the myogenic conversion of receptor-expressing C3H10T1/2 cells, evident by their expression of DsRed2 and conversion into syncytial cells (**Figure 5e-f**). Together with those described above, these results show that fluorescein-based ligands can direct gene expression and cell activities depending on the composition and folding state of ECM constituent proteins.

## DISCUSSION

In this study, we leveraged exogenously administered ‘adaptor’ compounds to devise new ways to regulate SynNotch activity via bio-orthogonal and photo-chemical reactions and in response to the folding state of natively-derived collagen-I proteins. To do so, we combined an optimized fluorescein-binding SynNotch with biocompatible ‘adaptors’ based on fluorescein- and OG448-conjugates. Using this approach, we demonstrated spatial, temporal, and dose-dependent control over *α*FITC(E2)-SynNotch signaling and its downstream outcomes. As a utility of these new methods, we exploited them to direct the myogenic conversion of multipotent fibroblasts into myoblasts using diverse triggers, including bio-orthogonal bond formation, photo-patterned ligands, and unfolded collagen—a disease biomarker associated with osteoarthritis and other pathological conditions^57^.

The FDA-approved and bio-compatible nature of fluorescein may facilitate facile extensions of this approach to *in vivo* settings, where fluorescein- and OG488-based adaptors could activate and confine therapeutic cell activities to targeted sites within the body. For example, TCO-functionalized materials could be combined with exogenous Tz-5-fluorescein administration to generate signaling-competent ligands within disease-relevant sites containing implanted biomaterial scaffolds. Additionally, TCO-containing nanoparticles could be directed to tumor sites via enhanced permeability and retention (EPR)^64^ and subsequently modified via Tz-5-fluorescein dosing to confine *α*FITC(E2)-SynNotch agonists to malignant sites *in vivo*. While our studies exploited a Tz-containing ligand, compounds based on methyl-Tz handles are likely to behave similarly while having improved serum and storage stabilities^65^.

Using a bacterially derived CNA35 domain, we directed cells to sense folded collagen-I and enact dose-dependent responses based on gene transcription. Additionally, by anchoring CHP-fluorescein to unfolded collagen strands, we devised a synthetic signaling strategy mimicking that of natural ECM regulation, in which mechanical unfolding is used to reveal cryptically buried signals in ECM proteins^66,67^. While heat-denatured collagen-I was applied as a model in our studies, CHPs are reported to bind diverse forms of denatured and damaged collagens, including protease-digested strands and mechanical ruptured fibrils, which are hallmarks of conditions such as aging and cancer^68,69^. In future studies, our approach could be extended to enable the detection of diverse disease-specific ECM-based signatures (beyond denatured collagens), including post-translationally modified ECM components, cleavage-generated neo-epitopes, or alternatively-spliced disease-associated ECM protein isoforms^70,71^. The detection of such species could be facilitated through fluorescein-labeled binding peptides, bacterial proteins, or fluorescein-modified antibody conjugates.

While the fluorescein adaptor-based approach makes this system highly versatile, users should be aware of potential limitations when applying or adapting the techniques that are described herein. Firstly, heterobifunctional compounds are susceptible to the dose-mediated “hook” effect, in which treatment with adaptor concentrations in excess can result in the non-productive occupation of binding sites with reduced or without the desired ligand-receptor bridging. Thus, bifunctional bridging compounds should be screened across various doses to identify suitable concentrations, which may vary depending on each compound, their applications, or the media compositions in which they are tested. New agents should be evaluated to confirm their independent binding to target components (including sender and receiver cells, immobilized scaffolds, ECM proteins, etc.) before cell-signaling measurements. Second, while the vast majority of commercial fluorescein conjugates will be based on 5- and 6-linked isomer derivatives, dyes linked via alternative positions may not be recognized by the *α*FITC(E2) scFv. Thus, chemical structures for conjugated dyes should be inspected or requested from vendors before use.

Diverse fluorescein-based indicator dyes have been previously developed as sensing agents to report on an array of biological signals and activities, and many of these agents could be combined with our system to facilitate new synthetic sense-and-respond activities in the future. For example, fluorescein-based compounds with sensitivity to H_2_O_2_^72^, or those designed to detect the activity of proteases, esterases^73^, and glycosidases could be employed as micro-environmental sensors capable of directing gene expression activities in *α*FITC(E2)-SynNotch cells^74–76^. Through such approaches, existing dyes originally developed as fluorescent sensors could be leveraged in synthetic biology applications as inducers and actuators. Additionally, while we used CMNB-caged fluorescein to generate *α*FITC(E2)-SynNotch activity patterns in 2D, the approach could be readily extended to 3D applications, where techniques like multiphoton laser-scanning lithography^77^ could be leveraged to define gene expression patterns in hydrogel-based systems, organ-on-a-chip models^78^, and in the manufacturing lab-grown meat^79^. Lastly, while we exploited immobilized agents to induce signaling activation, approaches involving the regulated disassembly of tethered ligands (via chemical, optical, or enzyme-based triggers) could be exploited as strategies to mediate ligand inactivation. Overall, we developed new chemogenetic components based on fluorescein and a fluorescein-binding SynNotch, demonstrating their utility in controlling gene expression activities in mammalian cells in response to a diverse set of new chemical triggers and biological cues.

## ACKNOWLEDGEMENTS

This work was funded through NIH research grants R35GM128859 (to J.T.N.) and R01HL147585 (to C.S.C. and J.T.N.). J.C.T and R.E.M. were supported through a Cross-Disciplinary Fellowship from BUnano (Boston University Nanotechnology Innovation Center). J.C.T., Z.E.S., and D.C.S. received support through the Boston University training program in Quantitative Biology and Physiology (QBP, NIH grant T32GM008764). D.C.S. and A.M.M. were recipients of National Science Foundation Graduate Research Fellowships. Additional support for the work was provided through the Center for Multiscale & Translational Mechanobiology (CMTM, Boston University) and the Multicellular Design Program (MDP, Boston University).

## AUTHOR CONTRIBUTIONS

All authors contributed to the design of experiments and edited the manuscript. J.C.T., C.J.K, A.M.M, R.E.M., Z.E.S., J.B.M., D.C.S.., and J.T.N. executed experiments.

