## Supplementary Information for "Fluorescein-Based SynNotch Adaptors for Regulating Gene Expression Responses to Diverse Extracellular Cues"

**Supplementary Movie 1. Time-lapse imaging of H2B-mCherry expression in HEK293-FT $\alpha$FITC(E2)-SynNotch reporter cells in response to Tz-TCO ligation-mediated activation.** HEK293-FT reporter cells (UAS:H2B-mCherry) expressing myc-$\alpha$FITC(E2)-LaG16-SynNotch-T2A-BFP were grown in microwells containing immobilized TCO functional handles. Cells were treated with Tz-5-fluorescein at the indicated concentrations at t = 0. Live cell recordings of reporter expression levels were captured at 0.5-hour intervals over 24 hours. Top row: H2B-mCherry is depicted in red, T2A-BFP in blue, overlaid with transmitted light. Bottom row: H2B-mCherry intensities depicted in grayscale.

**Supplementary Movie 2. Time-lapse imaging of DsRed2 expression in U2OS $\alpha$FITC(E2)-SynNotch reporter cells in response to Tz-TCO ligation-mediated activation.** U2OS reporter cells (UAS:DsRed2) expressing myc-$\alpha$FITC(E2)-LaG16-SynNotch-T2A-BFP were grown in microwells containing immobilized TCO functional handles. Cells were treated with Tz-5-fluorescein at the indicated concentrations at t = 0. Live cell recordings of reporter levels were captured at 0.5-hour intervals over 24 hours. Top row: DsRed2 is depicted in red, T2A-BFP in blue, overlaid with transmitted light. Bottom row: DsRed2 intensities depicted in grayscale.

**Supplementary Movie 3. Live cell imaging of reporter activity in response to BSA-PC-5-fluorescein photo-uncaging.** Live cell imaging of HEK293-FT reporter cells (UAS:H2B-mCherry) expressing a Gal4-VP64-containing $\alpha$FITC(E2)-SynNotch receptor. Cells were grown on gridded imaging dishes coated with BSA-PC-5-fluorescein. At t = 0, BSA-PC-5-fluorescein was photo-uncaged via a 100-millisecond pulse illumination through a DAPI excitation filter using a 63x oil-immersion objective. Live cell time-lapse recording of reporter expression levels was then collected through a 40x oil immersion objective, with the photo-illuminated area positioned at the center of the captured field of view. Images were captured every hour for 24 hours. H2B-mCherry is depicted in red, overlaid with transmitted light.

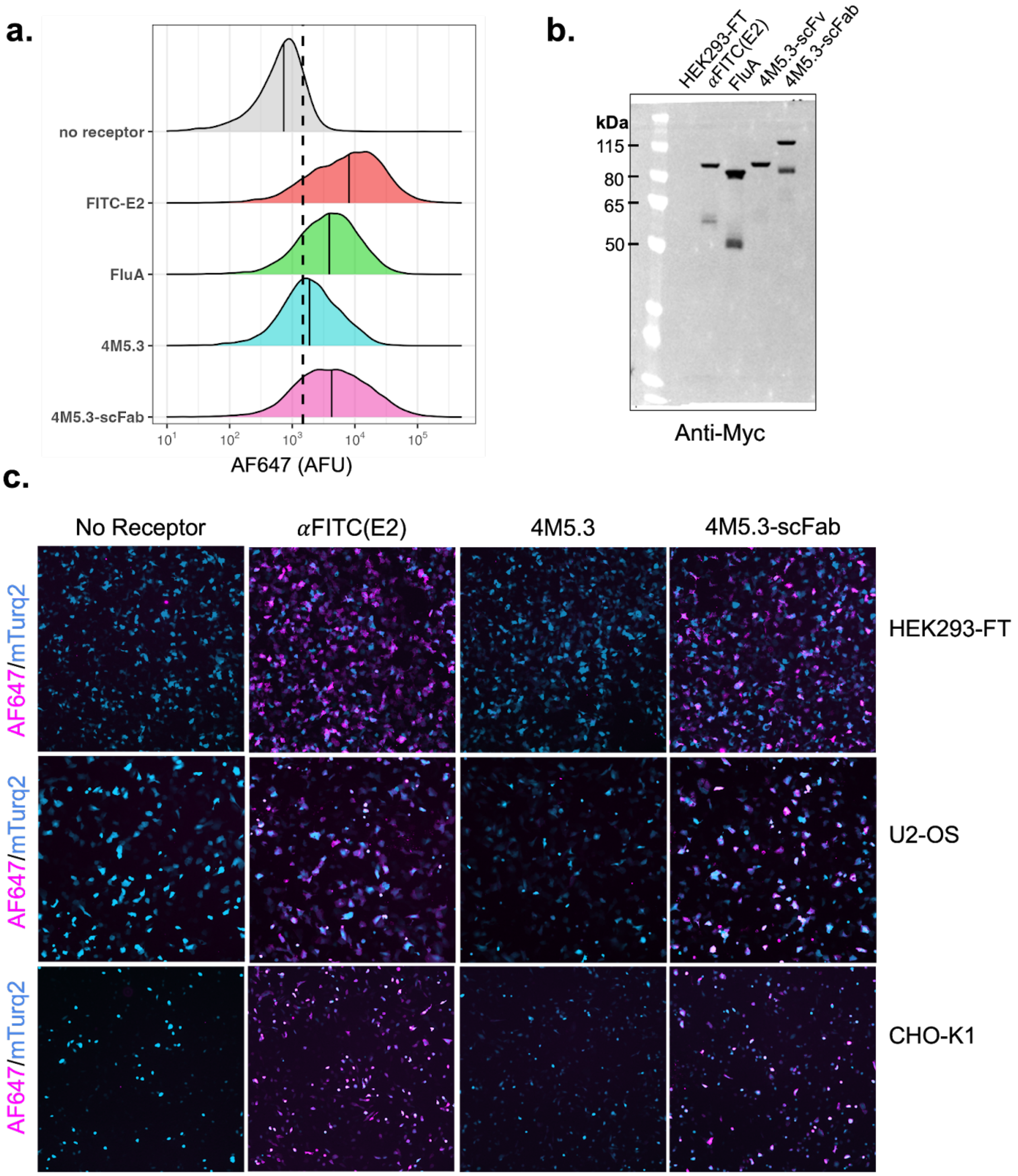


**Supplementary Figure 1. Surface expression of fluorescein-binding SynNotch receptors. (a)** HEK293-FT cells transfected with plasmid DNAs encoding the indicated receptors in combination with a construct encoding mTurquoise2 as a co-transfection marker. Cells were stained the following day using an anti-myc-AF647 conjugate to label surface-localized receptor copies. Levels of the immunolabeled receptors were quantified via flow cytometry with gating for co-transfected (mTurquoise2 positive) cells. Representative traces are shown. Each trace represents an analysis of > 10,000 cells, which were transfected one day before analysis using 100 ng receptor-encoding plasmid and 10 ng pcDNA3-mTurquoise2. Black lines denote median AF647 intensities; the dashed black line indicates the AF647 threshold fluorescence values calculated from non-transfected HEK293-FT cells. **(b)** Immunoblotting shows that 4M5.3-scFab SynNotch receptors are processed into a noncovalent heterodimer via furin cleavage; lower bands represent myc-tagged N-terminal furin cleavage products, and higher bands represent full-length polypeptides. **(C)** Immunofluorescence images of non-permeabilized cells following staining with anti-myc-AF647. Images of cells expressing the indicated receptors and cotransfected with a mTurquoise2 encoding plasmid are shown. Analyses were performed using transfected HEK293-FT, U2OS, and CHO-K1 cells.

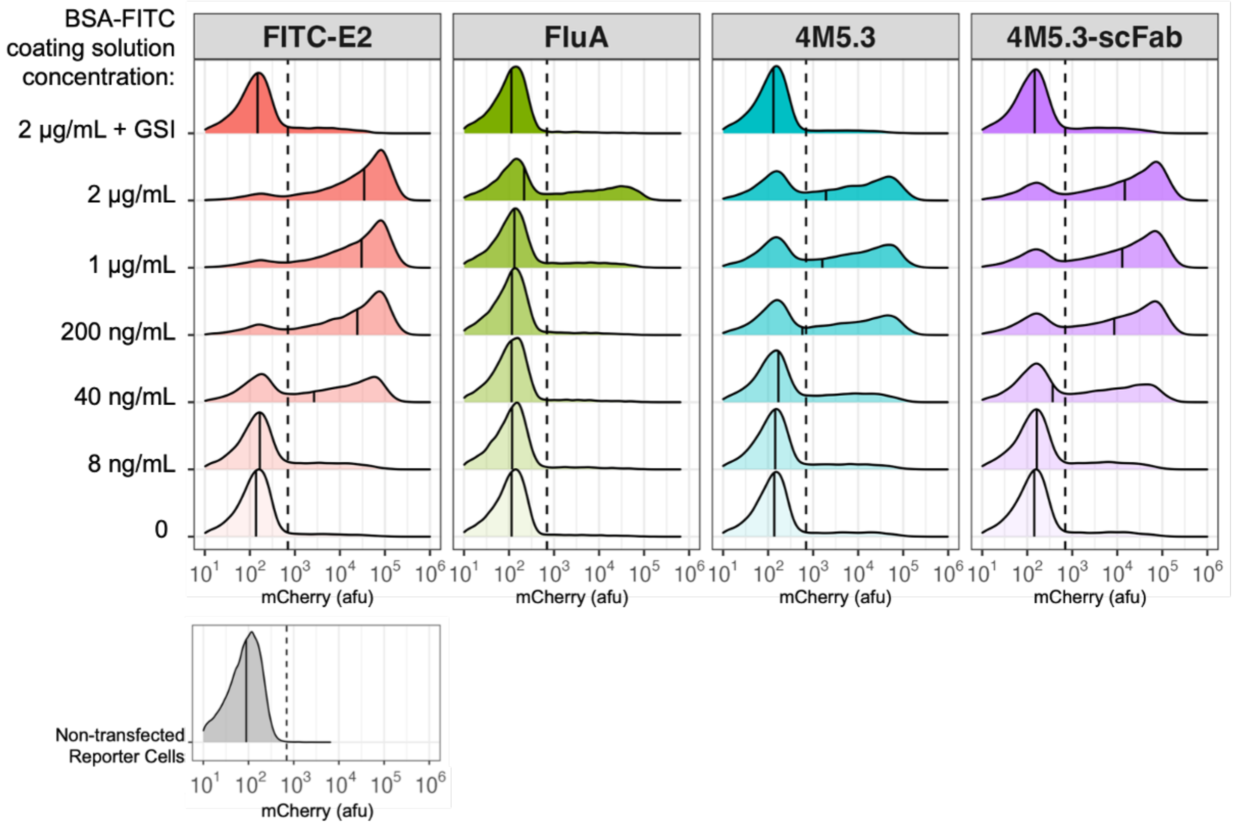


**Supplementary Figure 2. Dose-dependent signaling responses to immobilized BSA-5-FITC.** HEK293-FT (UAS-H2B-mCherry) reporter cells were transfected with DNA plasmids encoding the indicated receptors in combination with pcDNA3-mTurqoise2 as a co-transfection marker. Microwells were coated using BSA-5-FITC solutions at the indicated concentrations before adding transfected cells. Reporter H2B-mCherry levels were quantified via flow cytometry following overnight growth in coated microwells. Non-transfected reporter cells grown without ligand were analyzed as a control. Normalized traces of three independent transfections with gating based on the expression of the co-transfection marker are displayed (n = 3, > 5,000 cells per replicate). Ligand-treated cells grown with gamma-secretase inhibitor DAPT (5 µM, GSI) are also shown. Solid black lines denote median mCherry intensities; dashed black lines depict the fluorescence thresholds used to identify H2B-mCherry+ cells, as set based on analysis of non-transfected HEK293-FT reporter control cells.

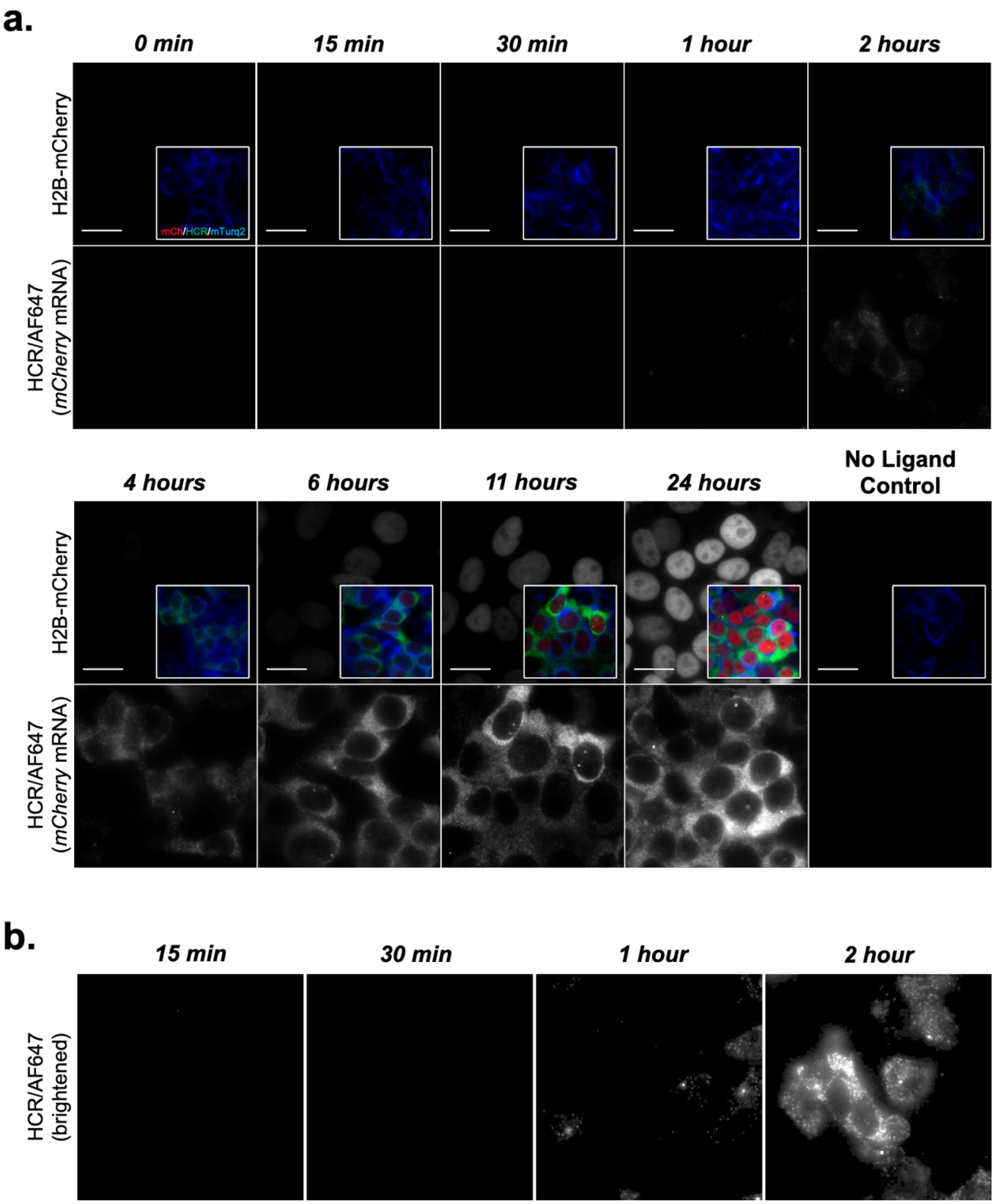


**Supplementary Figure 3. Reporter mRNA detection after GSI removal.** HEK293-FT (UAS-H2B-mCherry) reporter cells stably expressing a mTurq2-fused receptor were grown for 24 hours in coverslip bottom imaging wells coated with gelatin-FITC. Cells were added to the microwells in the presence of DAPT (5 μM) to block gamma-secretase-mediated ICD cleavage temporarily. Release from GS inhibition was facilitated by removing the DAPT-containing media, followed by rinsing three times before returning cells for growth in pre-warmed DAPT-free media. Cells were allowed to process ligand-activated receptors by gamma-secretase cleavage for the indicated times before fixation using 4% paraformaldehyde. The fixed specimens were then subjected to fluorescence *in situ* hybridization (FISH) analysis using Hybridization Chain Reaction (HCR) probes against *mCherry* mRNAs (see Methods). Scale bars = 25 μm.

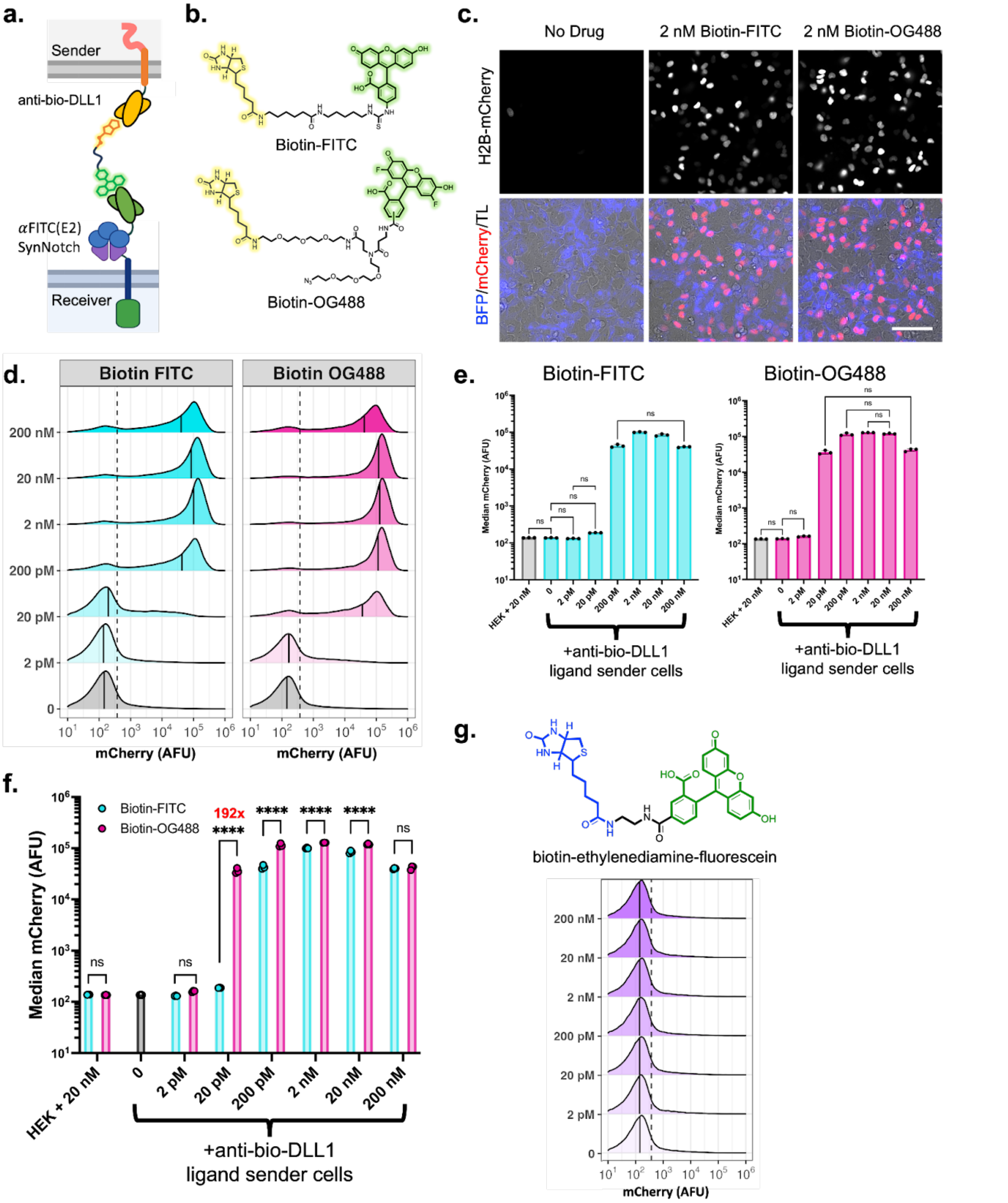


**Supplementary Figure 4. Comparison of bifunctional bridges containing either fluorescein or OG488 linked to biotin handles.** (**a**) Schematic depicting the bifunctional bridging strategy for $\alpha$FITC(E2)-SynNotch trans-activation by "sender" cells expressing the biotin-binding ligand (anti-bio-SNAP-TMD-DLL1). (**b**) Structures of the fluorescein- and OG488-containing biotin conjugates (biotin-FITC and biotin-OG488, respectively). (**c**) Fluorescence images of HEK293-FT receiver cells expressing $\alpha$FITC(E2)-LaG16-SynNotch-T2A-BFP in coculture with stable HEK293-FT sender cells expressing anti-bio-SNAP-TMD-DLL1. H2B-mCherry in grayscale (top) or red (bottom); T2A-BFP emissions are depicted in blue (bottom). Representative images of untreated cocultures and those treated with biotin-FITC or biotin-OG488 at 2 nM concentrations are shown. Images were captured at 24 hours post-drug treatment. Scale bar = 100 μm (**d**) Traces represent normalized densities of three independent cocultures (n = 3, >5,000 cells analyzed per replicate) of $\alpha$FITC(E2)-SynNotch-mTurq2 receivers grown with anti-bio-SNAP-TMD-DLL1 senders and treated with varying biotin-FITC and biotin-OG488. Receiver cells were identified by gating mTurq2+ cells. The dashed black line indicates the threshold value for defining H2B-mCherry+ cells. Thresholds for both fluorescent proteins were defined based on an analysis of HEK293-FT controls. The solid black line denotes median H2B-mCherry fluorescence intensities. The 0 nM bridging molecule concentration control is the same cell population, shown with alignment to drug-treated traces for reference and comparison. (**e**) Median receiver cell reporter intensities (H2B-mCherry) from (d) with either HEK293-FT (control) or anti-bio-SNAP-TMD-DLL1 sender cells at indicated concentrations of biotin-FITC and biotin-OG488. Co-cultures containing control senders (HEK293-FT) were grown with bridging compounds at 20 nM concentrations. Displayed values were analyzed using one-way ANOVA. All unlabeled comparisons in coculture conditions of various bridging concentrations with anti-bio-DLL1 senders versus 0 nM concentration have $P < 0.0001$; labeled NS, $P>0.05$. (**f**) Displayed values from (d) were analyzed using two-way ANOVA (bridging molecule and concentration). labeled NS, $P>0.05$, *$P < 0.05$, **$P<0.01$, ***$P<0.001$, ****$P<0.0001$. (**g**) structure of the inactive (short-linker containing) bridging agent biotin-ethylenediamine-fluorescein and dose-dependent analyses of the compound using cocultured cells in the same manner as in (d-f). Biotin-ethylenediamine-fluorescein failed to mediate receiver cell trans-activation at the tested concentrations.

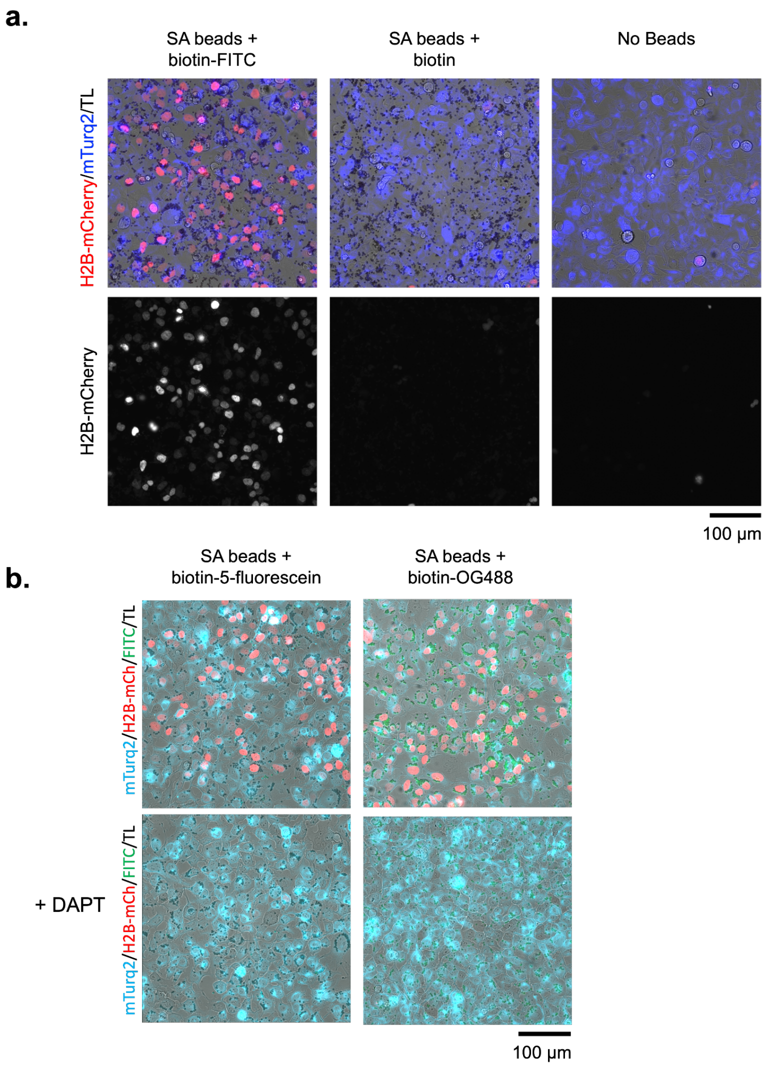


**Supplementary Figure 5. $\alpha$FITC(E2)-SynNotch activation by dye-decorated microbeads.** A stable line of HEK293-FT (UAS:H2B-mCherry) cells expressing $\alpha$FITC(E2)-SynNotch-mTurq2 was incubated with streptavidin (SA)-coated microbeads labeled with (**a**) biotin as a free acid (biotin) or biotin-FITC, and **(b)** biotin-FITC (a.k.a., biotin-5-fluorescein; same as in (a)) or biotin-OG488. Co-treatment of cells with DAPT (5 µM) blocked ligand-mediated activities. Cells were imaged 24 hours after applying beads to cells. H2B-mCherry is depicted in red (both), and mTurq2 is shown in blue for (a) and in cyan for (b). For (b), FITC and OG488 were imaged under the same excitation and emission settings and are rendered in green. The increased emission from biotin-OG488 beads is attributed to OG488's increased stability within the physiological pH range.

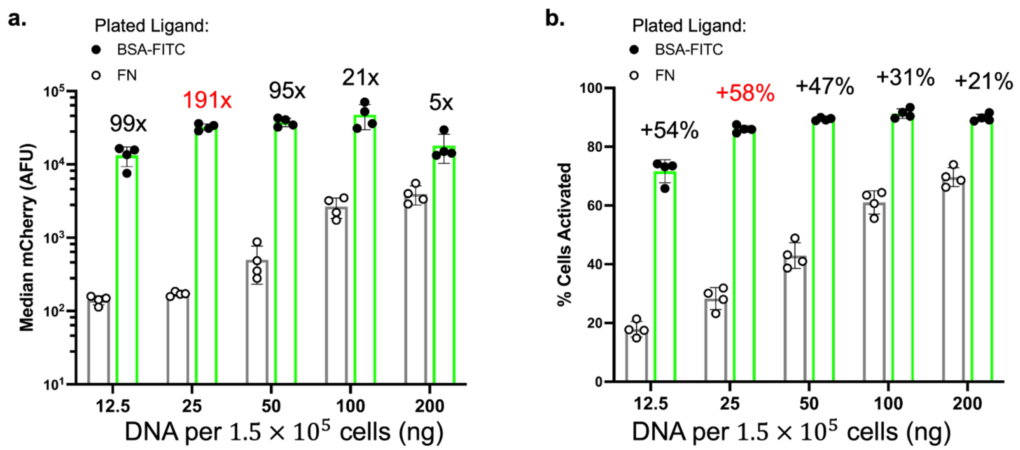


**Supplementary Figure 6. Optimization of transient receptor expression levels.** Ligand-induced and background activities were analyzed in HEK293-FT (UAS:H2B-mCherry) reporter cells transiently transfected with plasmid DNA encoding a receptor with an extracellular domain based on $\alpha$FITC(E2) and the LaG17 anti-GFP nanobody (pcDNA3-$\alpha$FITC(E2)-LaG17-SynNotch-Gal4VP64). Cells were transfected with the indicated amounts of pcDNA3-based plasmid. Transfected cells were analyzed by flow cytometry following overnight growth in fibronectin-coated (control) versus fibronectin and BSA-5-fluorescein-coated microwells (n = 4, > 5,000 cells per replicate). Gating was performed based on the co-expression of a mTurq2 co-transfection marker (expressed in cells via a co-transfected pcDNA3-mTurqoise2 plasmid). Gal4VP64-mediated expression levels of the H2B-mCherry reporter were then quantified 24 hours after plating. **(a)** Median H2B-mCherry fluorescence intensities in mTurq2+ cells are depicted. Fold changes are displayed as ratios between the means of median fluorescence values for ligand-treated versus untreated conditions. (**b**) Reporter activity levels are shown as percentages of H2B-mCherry+ cells within analyzed mTurq2+ populations. H2B-mCherry+ cells are defined as those containing emission levels above an intensity threshold based on analysis of non-transfected HEK293-FT cells (> top 99th percentile). The indicated DNA amounts correspond to levels of receptor-encoding DNA used to prepare the Lipofectamine 3000 transfection mixtures provided to cells.

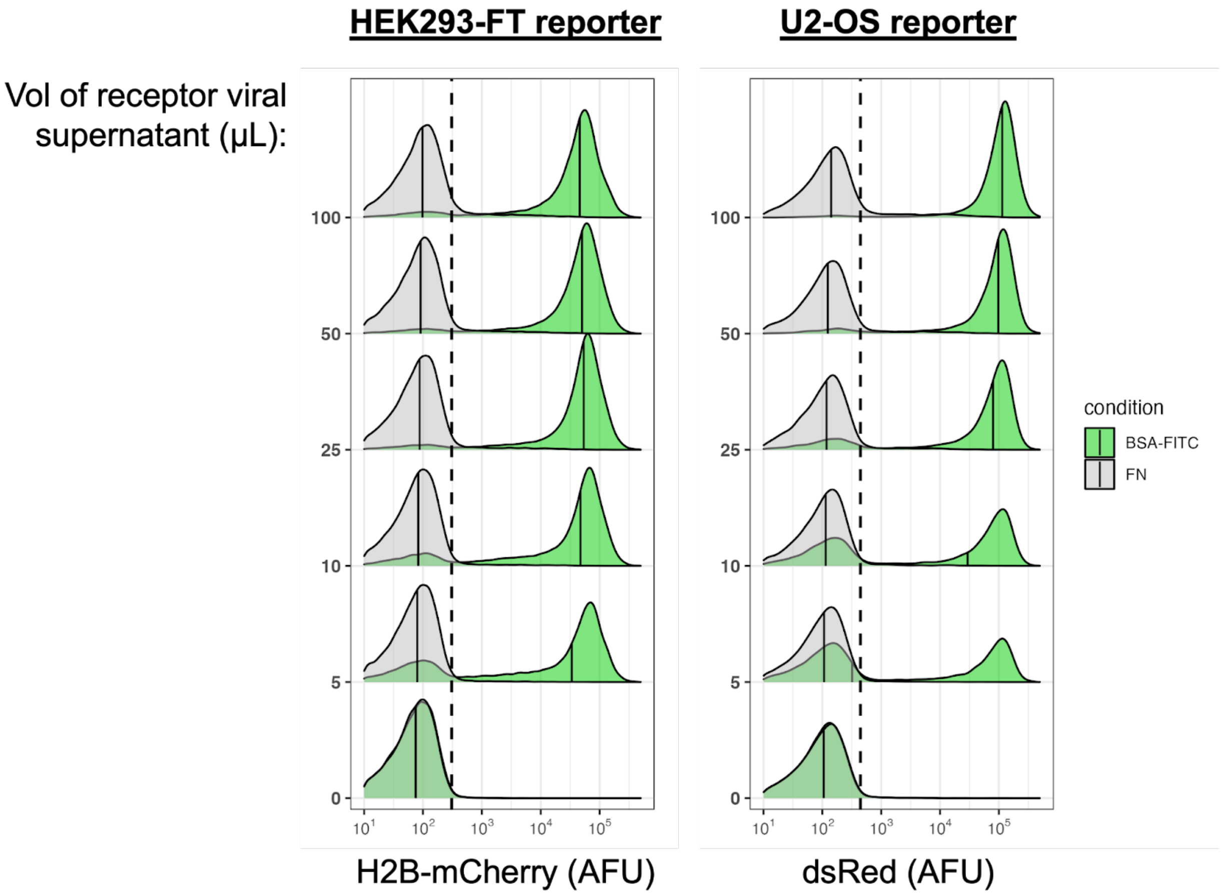


**Supplementary Figure 7. Lentiviral transduced receptor in HEK and U2OS reporter cells.** A lentiviral construct encoding $\alpha$FITC(E2)-SynNotch-Gal4VP64 from an SFFV promoter was used to generate viral particles and transduce cells. Virus-containing supernatants were applied to HEK293-FT (UAS:H2B-mCherry, left) or U2OS (UAS:DsRed2, right) reporter cells at the indicated supernatant volumes. Cells were analyzed by flow cytometry following 36 hours of growth in fibronectin-coated (control) versus fibronectin and BSA-5-FITC-coated microwells. Gating was performed based on the co-expression of tagBFP2 (via co-transduction of 10 µL viral supernatant volumes harvested from cells producing particles packaged with pHR-SFFV-T2A-tagBFP2). Traces represent normalized densities (>10,000 cells). Black lines denote median reporter fluorescence intensities; the dashed lines indicate the reporter intensity thresholds used to define activated cells, as determined using non-transduced control reporter cells (> top 99th percentile).

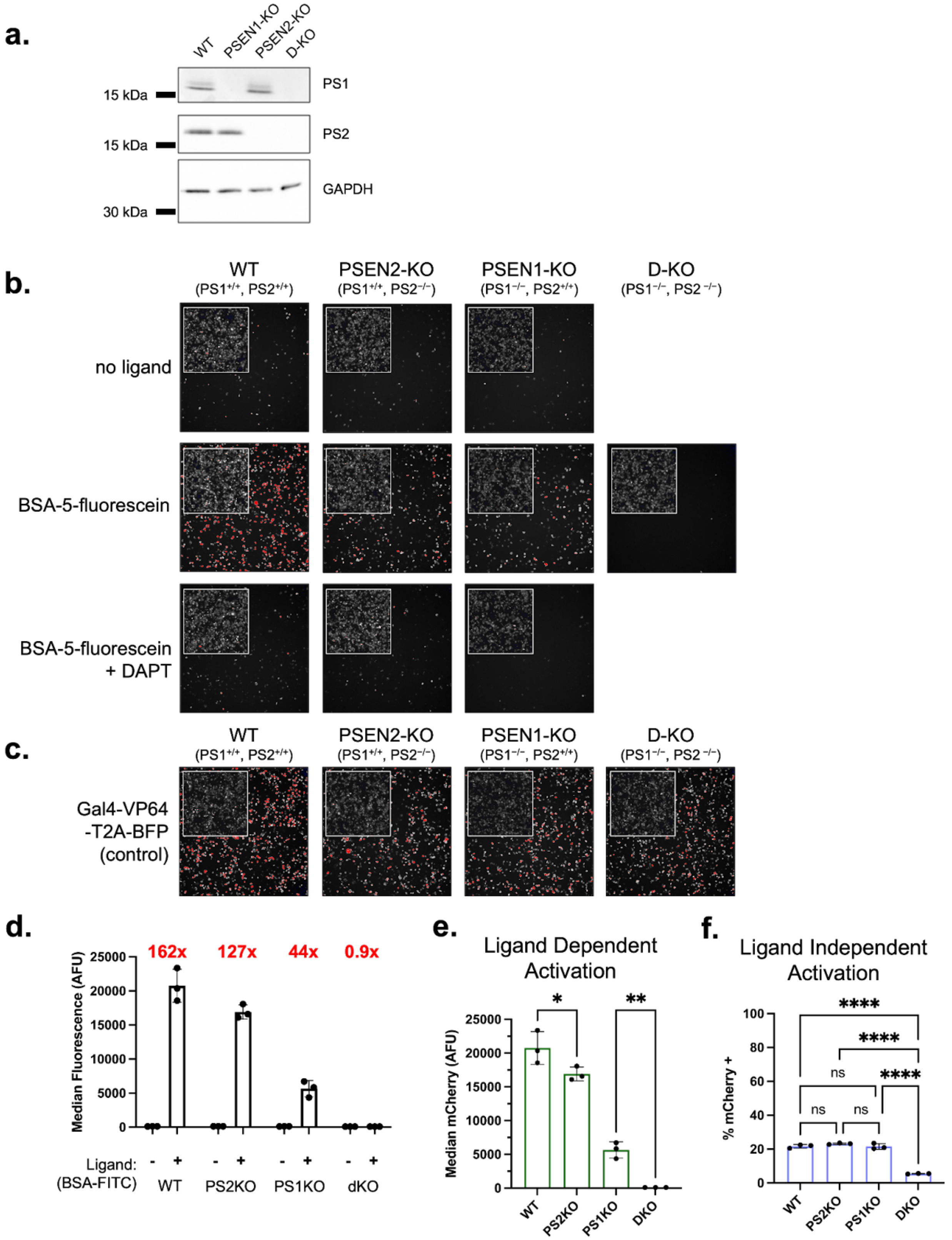


**Supplementary Figure 8. Functional characterization of presenilin-1 and presenilin-2 knockout (KO) in $\alpha$FITC(E2)-SynNotch expressing HEK293-FT cells.** (**a**) Immunoblot of reporter HEK293-FT UAS:H2B-mCherry KO cell lysates confirming successful KO of PS1, PS2, and double KO of PS1 and PS2. (**b**) H2B-mCherry activation in PS KO lines transfected with $\alpha$FITC(E2)-SynNotch-T2A-BFP and cultured on fibronectin- (no ligand) and BSA-5-fluorescein-coated microwells. DAPT (5 µM) treatment was used to block ligand-mediated signals. Insets represent emissions from the expression of a T2A-BFP co-expression marker. (**c**) Plasmid encoding a positive control transcription factor based on GAL4-VP64-T2A-BFP confirmed the maintained sensitivity of the KO reporter lines to Gal4-mediated activation of the UAS:H2B-mCherry reporter gene. (**d-f**) H2B-mCherry activation quantified by flow cytometry from three independent transfections gated for BFP+ from non-transfected control cells (n = 3, > 5,000 cells assessed per replicate). Fold-change values indicated in (d) represent ratios between means of median fluorescence values. (**e**) Activation on plated BSA-5-fluorescein was analyzed by one-way ANOVA. All unlabelled comparisons have P < 0.0001; labeled NS: P > 0.05, *: P <0.05, **: P <0.005, otherwise. (**f**) H2B-mCherry expressing fractions of transfected and ligand-untreated cell represented as percentages of BFP+ populations. H2B-mCherry-expressing cells were defined as those exhibiting emission intensities beyond a threshold set based on the analysis of non-transfected control reporter cells. Analyzed by one-way ANOVA. dKO was found to decrease ligand-independent activation significantly. NS: P > 0.05, ****: P < 0.00001.

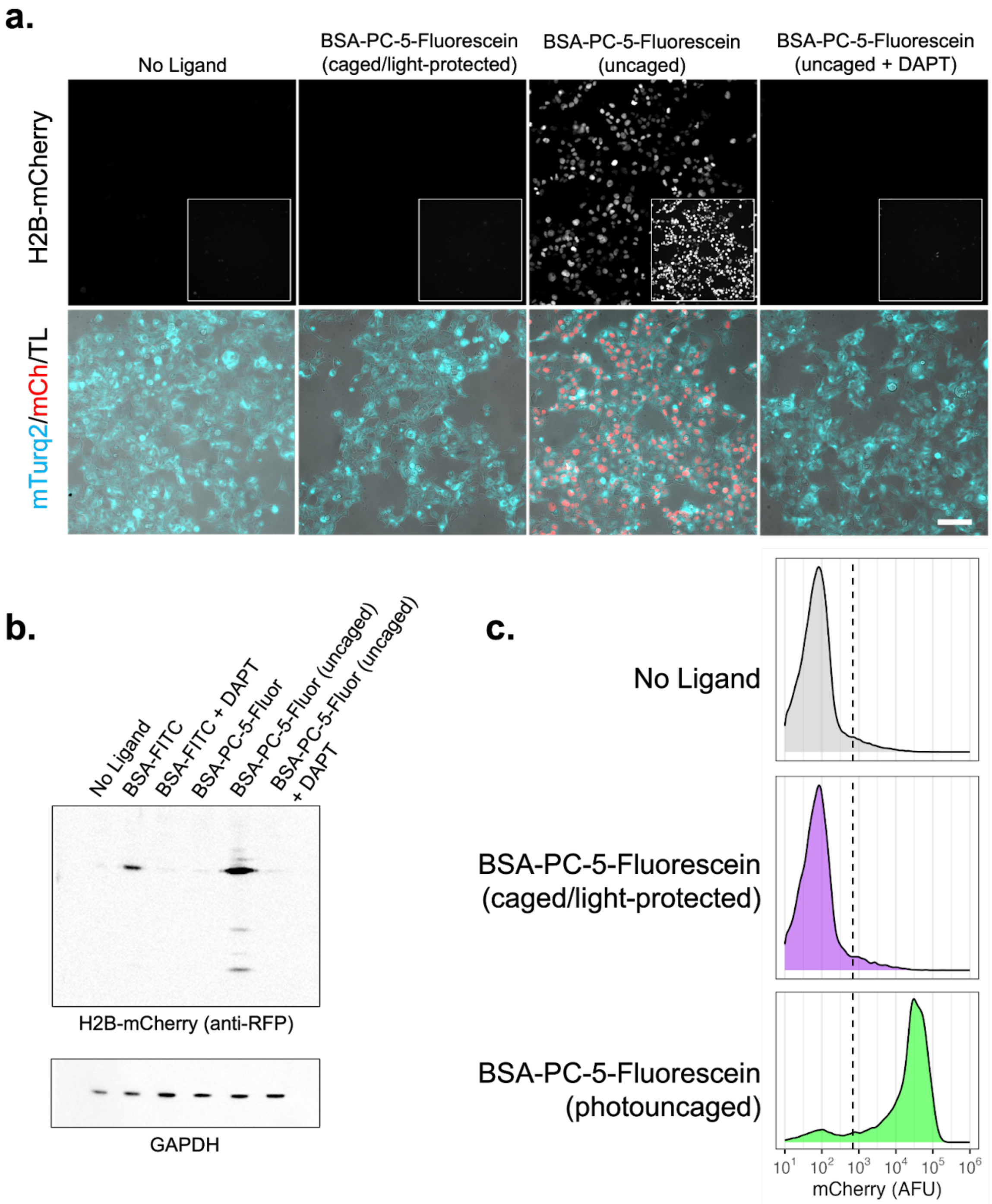


**Supplementary Figure 9. BSA-PC-5-fluorescein is a light-dependent SynNotch ligand.** Clonal HEK293-FT (UAS:H2B-mCherry) reporter cells expressing $\alpha$FITC(E2)-SynNotch-mTurq2 were grown in microwells containing caged (light protected) or uncaged BSA-PC-5-fluorescein. **(a)** Fluorescence imagines of reporter activities from cells grown under the indicated conditions; H2B-mCherry is depicted in grayscale (top) and red (bottom); mTurq2 is shown in cyan (bottom). Scale bar = 100 µm. Insets represent H2B-mCherry emissions depicted in grayscale with increased digital contrast. **(b)** Analysis of reporter H2B-mCherry expression by immunoblot detection. **(c)** Flow cytometry measurement of cells grown under the indicated conditions. The dashed black line represents the intensity threshold used to define H2B-mCherry+ cells, as set based on the analysis of control HEK293-FT cells. The analyses in this figure were done using caged and uncaged BSA-PC-5-fluorescein; uncaged fluorescein was generated by subjecting BSA-PC-5-fluorescein-coated glass bottom culture dishes to transillumination with a handheld UV lamp for 1 minute.

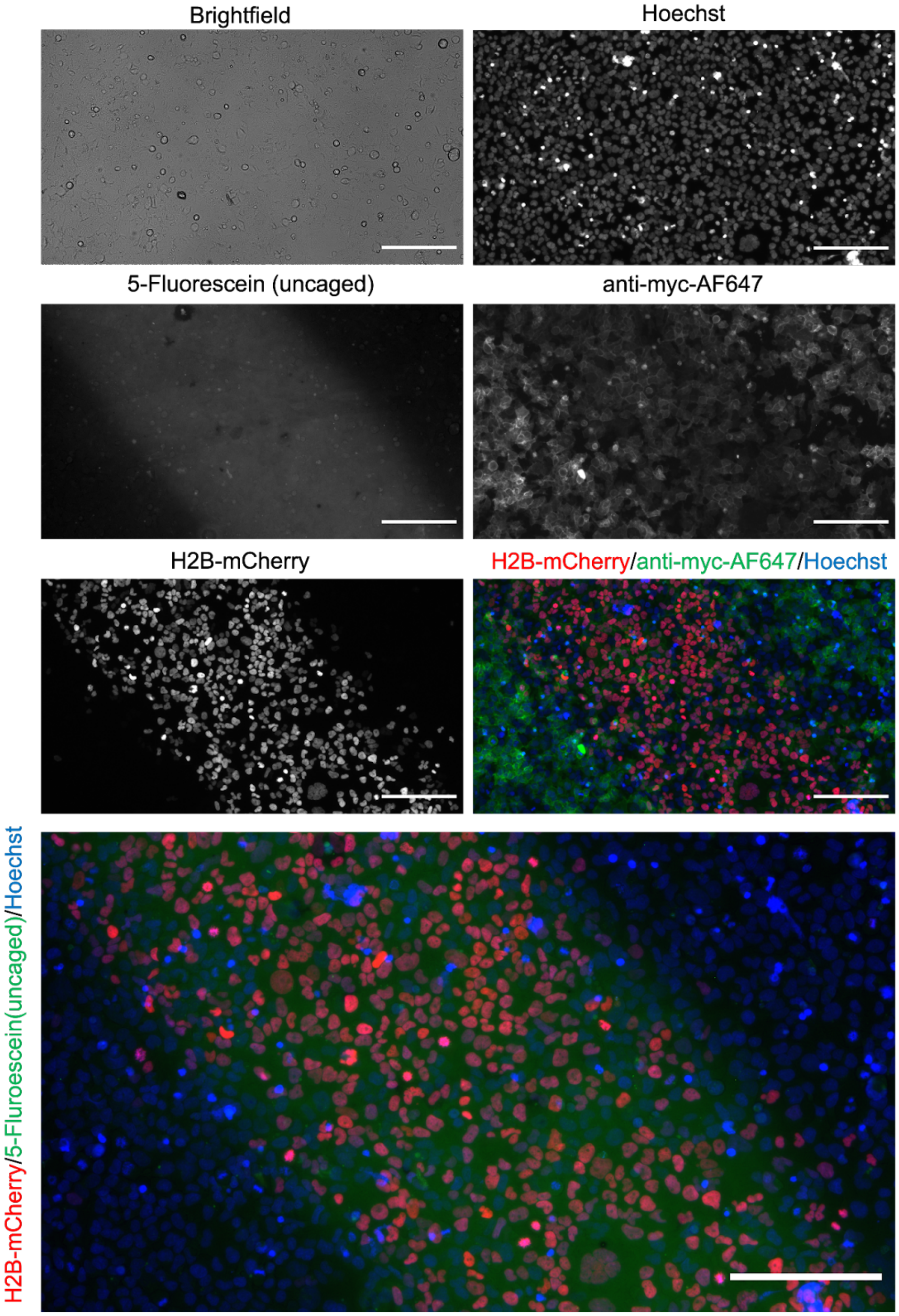


**Supplementary Figure 10. Spatial Control of $\alpha$FITC(E2)-SynNotch activation on photo-uncaged BSA-PC-5-Fluorescein substrate.** Epifluorescence images of a stable HEK293-FT cell line expressing the UAS:H2B-mCherry reporter and an $\alpha$FITC(E2)-SynNotch receptor plated on a dish that was coated with BSA-PC-5-Fluorescein. The bottom of the culture dish was taped over with black electrical tape, with an exposed ~1 cm diagonal gap before seeding cells. Photo-uncaging was done by transilluminating the well with a hand-held UV lamp for < 1 min. H2B-mCherry expressing cells coincide with regions of fluorescence emission from photo-uncaged PC-5-fluorescein ligands, as detected through the FITC imaging channel. Staining with anti-myc-AF647 was used to detect surface localized receptors; receptor-expressing cells outside the photo-uncaged region remained in signaling quiescent states. Scale bar = 200 µm.

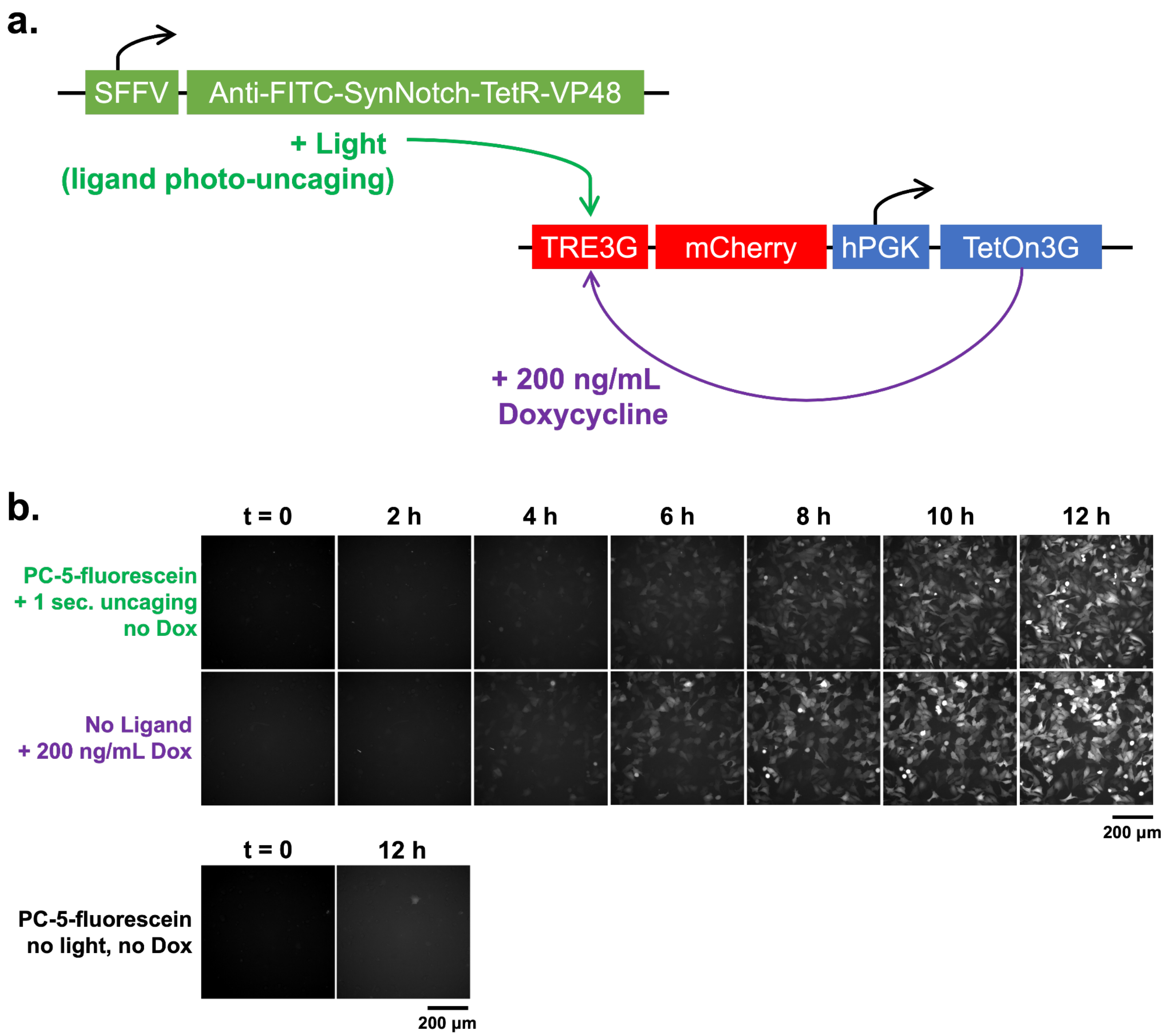


**Supplementary Figure 11. Photo-uncaging induced signaling compared to TetOn3G doxycycline induction. (a)** U2OS reporter cells (TRE3G:mCherry, hPGK:TetOn3G) were transduced with pHR-SFFV-$\alpha$FITC(E2)-SynNotch-TetR-VP48 lentiviral particles and **(b)** plated on indicated condition in doxycycline/tetracycline-free media. Top: at t = 0, wells were illuminated through a DAPI excitation filter to facilitate the photo-uncaging of BSA-PC-5-fluorescein ligand. Bottom: at t = 0, cells were exchanged into media containing doxycycline at 200 ng/mL to induce the activity of a co-expressed TetOn3G transactivator protein. Following treatment, time-lapse images of mCherry levels were captured every 2 hours for 12 hours.

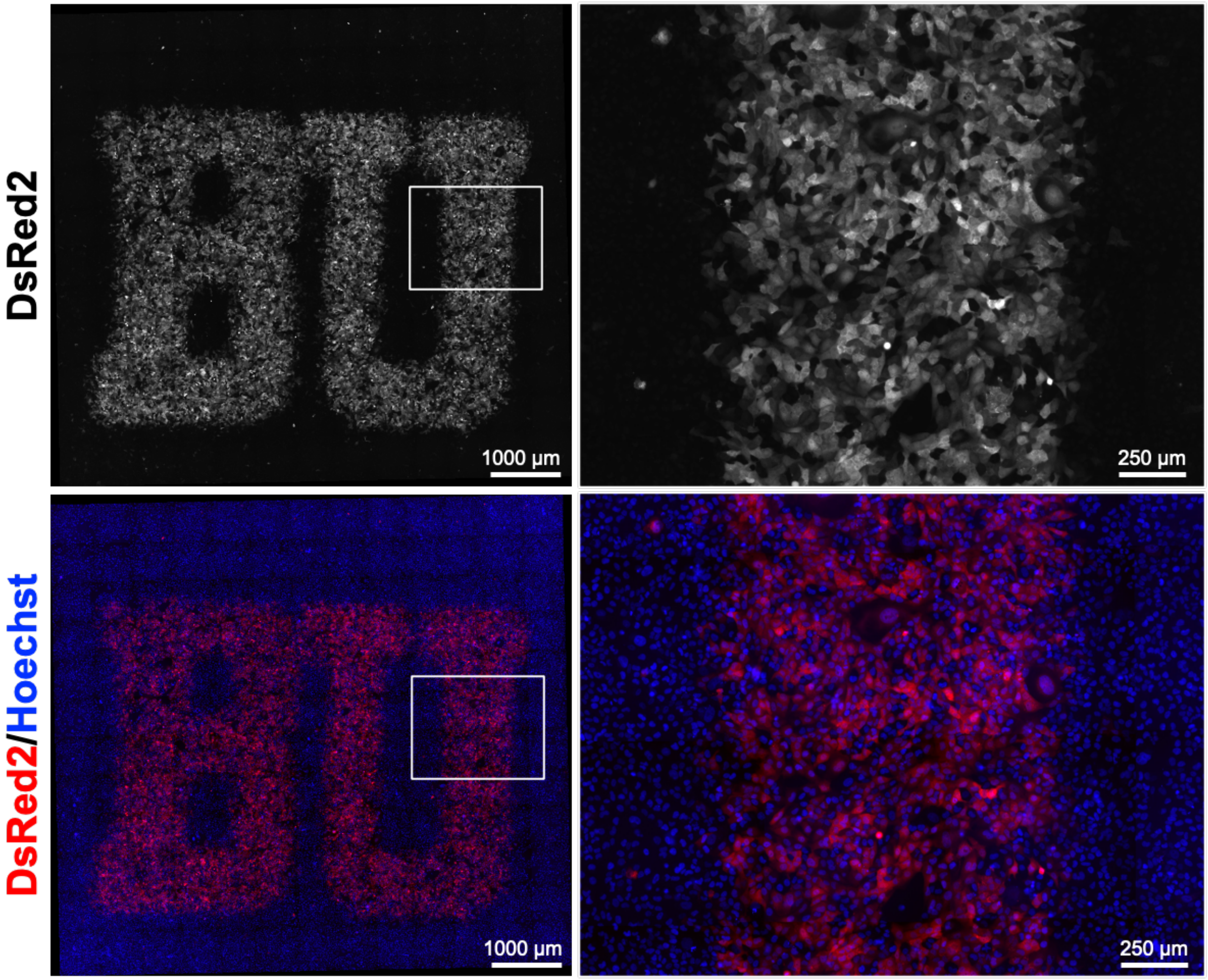


**Supplementary Figure 12. Spatial control of $\alpha$FITC(E2)-SynNotch-TetR-VP48 activation in C3H10T1/2 fibroblast containing a TRE-DsRed2 reporter.** C3H10T1/2 fibroblasts containing an integrated reporter gene based on TRE:DsRed2 (without p65-MyoD) were transduced viral constructs passed on pHR-SFFV-$\alpha$FITC(E2)-SynNotch-TetR-VP48. At 48 hours post-transduction, cells were seeded into a microwell containing immobilized and photo-patterned BSA-PC-5-fluorescein, similar to Figure 5D. Cells were imaged 3 days after plating.
