## Supplementary material for "Fluorescein-Based SynNotch Adaptors for Regulating Gene Expression Responses to Diverse Extracellular Cues": Methods

### REAGENTS AND MATERIALS

#### *DNA constructs*

*Receptors:*
pcDNA3-myc-$\alpha$FITC(E2)-SynNotch-Gal4-VP64
pcDNA3-myc-FluA-SynNotch-Gal4-VP64
pcDNA3-myc-4M5.3-scFv-SynNotch-Gal4-VP64
pcDNA3-myc-4M5.3-scFab-SynNotch-Gal4-VP64
pcDNA3-myc-LaG17-$\alpha$FITC(E2)-SynNotch-Gal4-VP64 (Sloas et al. 2023)
pHR-SFFV-myc-$\alpha$FITC(E2)-SynNotch-Gal4-VP64
pHR-SFFV-myc-$\alpha$FITC(E2)-SynNotch-TetR-VP48
pLV-EF1a-myc-$\alpha$FITC(E2)-LaG16-SynNotch-Gal4-VP64-T2A-BFP (Sloas et al. 2023; AddGene #194929)

*Lentivirus-Based Reporter Constructs:*
pLV-TRE-dsRed-Express2 (Kabadi et al. 2015; AddGene #60623)
pLV-TRE3G-mCherry, hPGK:TetOn3G

*CRISPR/Cas9 Knockout Plasmids:*
pGuide-it-tdTomato-hPSEN1-sgRNA
pGuide-it-tdTomato-hPSEN2-sgRNA

*Co-Expression Markers:*
pcDNA3-mTurquoise2 (transient transfection)
pHR-SFFV-T2A-tagBFP2 (viral transduction)

#### *Mammalian cell lines*

HEK293-FT (ThermoFisher, R70007)
U2OS (Sigma-Aldrich, 92022711-1VL)
CHO-K1 (Sigma, 85051005-1VL)
C3H/10T1/2, Clone 8 (ATCC, CCL-226)

*Reporter Cell Lines:*
HEK293-FT, UAS:H2B-mCherry (Sloas et al. 2023)
U2OS, UAS:DsRed-Express2, hPGK:Puro (Sloas et al. 2023; AddGene #190801)
U2OS, TRE3G:mCherry, hPGK:TetOn3G-T2A-Puro

C3H/10T1/2, TRE:p65-MyoD-2A-DsRed-Express2, hPGK:Puro (Sloas et al. 2023)
C3H/10T1/2, TRE:DsRed-Express2, hPGK:Puro

*Receiver (Receptor-Expressing) Cell Lines:*
HEK293-FT,UAS:H2B-mCherry,EF1a-$\alpha$FITC(E2)-SynNotch-Gal4-VP64 (clone)
HEK293-FT,UAS:H2B-mCherry,EF1a-$\alpha$FITC(E2)-SynNotch-Gal4-VP64-mTurq2 (clone)
HEK293-FT,UAS:H2B-mCherry,EF1a-LaG16-$\alpha$FITC(E2)-SynNotch-Gal4-VP64-T2A-BFP (pool)
U2OS,UAS:DsRed-Express2,EF1a-LaG16-$\alpha$FITC(E2)-SynNotch-Gal4-VP64-T2A-BFP (clone)

*Sender (Ligand-Expressing) Cell Lines:*
HEK293-FT, anti-bio-SNAP-TMD-DLL1 (Sloas et al. 2023; McMahan and Ngo 2022)

*Culture Media*
DMEM/High glucose with sodium pyruvate; without L-glutamine (Cytiva, SH30285.01)
Characterized Fetal Bovine Serum (FBS), Canadian origin (Cytiva, SH30396.03)
Tet-Approved FBS (Clontech, 631106)
Glutamax Supplement (ThermoFisher, 35050061)
Non-Essential Amino Acids (NEAA) Solution, 100X (Cytiva, SH30238.01)
Penicillin-Streptomycin Solution, 100X (Sigma-Aldrich, P4333-20ML)
HEPES solution, 1 M (Cytiva, SH30237.01)

*Commercially Available Chromophore-Based Ligand Conjugate Proteins and Related*
Gelatin solution, 2% (Sigma-Aldrich, G1393-100ML)
Gelatin, FITC Conjugated (AnaSpec, AS-85145)
Gelatin-OregonGreen488 Conjugate (ThermoFisher, G13186; 5-isomer, per supplier)
Bovine Fibronectin Protein, Carrier Free (R&D Systems, 1030-FN-05M)
BSA-FITC conjugate (Sigma, A9771; 12 mol. 5-FITC per mol BSA for lot used, per supplier)
Collagen Hybridizing Peptide (CHP), 5-FAM conjugate (Advanced Biomatrix, 5264)
Collagen-I, 4.04 mg/ml solution (Corning, 354236)

*Commercially Available Reactive Dyes, Bioorthogonal Handles, And Related*
FAM azide, 5-isomer (Lumiprobe, B4130)
6-FAM-azide (AAT Bioquest, Catalog No. 133)
Tetrazine-5-FAM (Jena Bioscience, CLK-013-05; *note: 6-Methyl-Tetrazine-5-FAM (CLK-018-1) is expected to have increased stability and storage lifetimes*)
TCO-PEG4-biotin (BroadPharm, BP-23847)
TCO-PEG4-NHS Ester (ClickChemistryTools, A137-2; CAS: 1621096-79-4; MW: 514.57)
CMNB-Caged Carboxyfluorescein, succinimidyl ester (ThermoFisher, C20050; MW: 962.78)

DBCO-NHS ester (ClickChemistryTools, A133-25)
Halo-DBCO (Iris Biotech, RL-3670; CAS: 1808119-16-5)
Bovine Serum Albumin (BSA), Heat Shock, Protease-Free (bioWORLD, 22070038-1)
20X Borate Buffer Stock, pH 8.5 (ThermoFisher, 28341)
Slide-A-Lyzer G2 Dialysis Cassettes, 10K MWCO, 3 mL capacity (ThermoFisher, 87730)
Slide-A-Lyzer G2 Dialysis Cassettes, 2K MWCO, 3 mL capacity (ThermoFisher, 66203)

*<u>Commercially available biotin-conjugated fluorophores</u>*

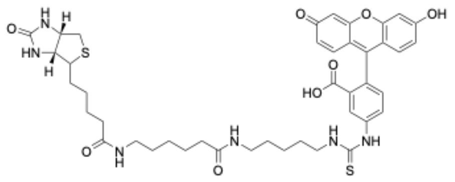

Biotin-FITC (Cayman Chemical, 25574; CAS: 134759-22-1; MW: 831.0)

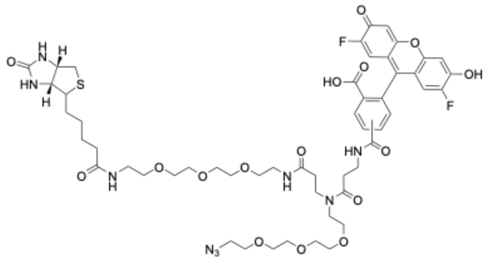

Biotin-OregonGreen488-azide (ClickChemistryTools, 1247-1; “Fluorescein Biotin Azide”; MW: 1156.44; 5/6 mixed isomer; now available from Vector Laboratories, CCT-1247)

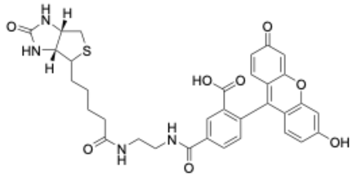

Biotin-4-fluorescein (ThermoFisher, B10570; “biotin-ethylenediamine-fluorescein”; CAS: 1032732-74-3; MW: 644.7)

*<u>Materials for protein expression and purification</u>*
OneShot BL21(DE3) Chemically Competent *E. coli* (ThermoFisher, C600003)
Ni-NTA Fast Start Kit (Qiagen, 30600)

IPTG (isopropyl-β-D-thiogalactopyranoside), Ultrapure (VWR, 97063-282)
2xYT broth powder, biotechnology grade, 1 kg (BioPioneer, CM2YT-1)

*Competent cells for Plasmid Preparation*
NEB Stable Competent *E. coli* (New England Biolabs, C3040H; for lentiviral vector plasmids)
DH5-Alpha Competent *E. coli* (BioPioneer, GACC-96)

*Materials*
8-well chambered cover Glass with #1.5 high-performance cover glass (Cellvis, C8-1.5H-N)
μ-Slide 18 Well Glass Bottom (ibidi, 81817)
Craftiff Clear Printable Transparency Film 8.5 x 11 in for Laser Printers (Amazon)
Non-Treated 96 Well Plates, polystyrene, sterile (Celltreat, 229596)

*Other Reagents and Materials*
Lipofectamine 3000 (ThermoFisher, L3000015)
SuperSignal West Pico PLUS Chemiluminescent Substrate (ThermoFisher, 34580)
Restore Western Blot Stripping Buffer (ThermoFisher, 21059)
PageRuler Prestained Protein Ladder, 10 to 180 kDa (ThermoFisher, 26617)
Dynabeads M-280 Streptavidin, 2.8-μm (ThermoFisher, 60210 or from the 65801D kit)
Biotinylated BSA (Sigma-Aldrich, A8549)
NeutrAvidin Protein (ThermoFisher, 31000)
Hoechst 33342, Trihydrochloride, Trihydrate, 10 mg/mL in water (ThermoFisher, H3570)
Verapamil (MedChemExpress, CP-16533-1)
JaneliaFluor-646 Hoechst (gift from L. Lavis, JaneliaFarm)
DAPT (gamma-secretase Inhibitor IX), 10 mM stock in DMSO (MedChemExpress, HY-13027)
Immunofluorescence Blocking Buffer (Cell Signaling Technologies, #12411)
GelCode Blue Stain Reagent (ThermoFisher, 24592)
Salmon sperm DNA (R&D Systems, 9610-5-D)
Custom HCR probe kit (Molecular Instruments)
16% Formaldehyde (w/v), methanol-free (ThermoFisher, 28906)
Low-Intensity UV Lamp, 110 V (Electron Microscopy Sciences 72422; VWR 100491-958)
10 kDa MWCO filters (66380, ThermoFisher)

*Primary Antibodies*

| *Antibody* | *Conditions (Immunofluorescence, IF; western blot, WB)* |
|---|---|
| Ms. monoclonal anti-Myc, clone 9E10, AlexaFluor647 (Santa Cruz Biotech., sc-40 AF647) | IF: 1:200, in culture media, 30 minutes at 37 °C in $CO_2$ incubator. |
| Ms. monoclonal anti-myosin heavy chain, clone MF20 (R&D Systems, MAB4470) | IF: 1:100 in antibody dilution buffer (PBS-T with 1% w/v BSA), overnight 4° C |
| Rb. polyclonal anti-RFP (Rockland, 600-401-379) | WB: 1:1000 in PBS-T, 1 hour at room temperature<br>WB: 1:1000 in blocking solution, 4° C overnight. |
| Rb. monoclonal anti-myc, clone 71D10 (Cell Signaling Technologies, 2278) | WB: 1:1000 in blocking buffer (PBS-T with 5% w/v nonfat dry milk), overnight 4° C |
| Direct-Blot anti-GAPDH-HRP (BioLegend, 607904) | WB: 1:4000 - 1:10,000 in PBS-T, 1 hour at room temperature |
| Rb. anti-Presenilin 1 (D39D1), mAb (Cell Signaling Technologies, 5643) | WB: 1:1000 in TBS-T with 5% w/v BSA, 4° C overnight after blocking membrane in PBS-T 5% w/v milk |
| Rb. anti-Presenilin 2 (D20G3), mAb (Cell Signaling Technologies, 9979) | WB: 1:1000 in TBS-T with 5% w/v BSA, 4° C overnight after blocking membrane in PBS-T 5% w/v milk |

*Secondary Antibodies*

| *Antibody* | *Conditions (Immunofluorescence, IF; western blot, WB)* |
|---|---|
| Goat Anti-Rabbit IgG (H + L)-HRP (Bio-Rad, 1706515) | WB: 1:3000 in PBS-T, 1 hour at room temperature |
| Rb. anti-mouse IgG (H+L) secondary, AlexaFluor647 (ThermoFisher, A21239) | IF: 1:1000 in antibody dilution buffer (PBS-T with BSA at 1% w/v), 1 hour at room temperature |

**Mammalian cell culture**

Mammalian cell lines were cultured in a humidified incubator maintained at 37° C with 5% $CO_2$. Cell media based on DMEM was supplemented with 10% FBS, 1x GlutaMAX, 1x non-essential amino acids, and 1x Penicillin-Streptomycin. For experiments involving cells containing doxycycline-inducible reporter genes, mediate containing 10% Tet-Approved (or otherwise-doxycycline-free) FBS was used in place of regular FBS (see Materials).

**Flow cytometry**

Cells were analyzed using an Attune NxT flow cytometer v2.6 and v3.1 flow cytometry software (ThermoFisher) as in (Sloas et al. 2023). Live cells were identified, setting FSC-A and SSC-A thresholds, and singlets were identified by a polygon gate set by FSC-A versus FSC-H, respectively. Positively transfected or transduced cells were identified based on the expression of a co-transfection marker or that of a co-translated T2A-fluorescent protein; positive cells were identified based on gates set using signals from control (non-expressing) cells (> 99th percentile). To improve the detection of cells expressing both a receptor of interest and co-transfection marker in transiently transfected cells, positively transfected cells were further gated for those bearing marker fluorescence emission at levels above the median of all transfected cells. Flow cytometry data were analyzed and quantified using the open-source ggCyto software (version 1.27.1).

**Data collection and analysis software**

Fluorescence images were collected using the Zen 2.3 Pro (Blue Edition) imaging software and analyzed in ImageJ v2.0.0. Western blots were collected and analyzed using the QuantityOne(4.5.2) immunoblotting software or the iBright Imaging System software v1.4.0. The data shown in the figures are representative examples of results that were repeated in at least two independent experiments. Statistical analyses were performed using the GraphPad Prism v9.0.0 software.

**Live time-lapse Imaging**

For live time-lapse imaging, cells were cultured in FluoroBrite DMEM supplemented with 5% FBS, 1X GlutaMAX, 1X non-essential amino acids, and containing 20 mM HEPES (pH 7.4). Imaging was performed in media containing JaneliaFluor-646 Hoechst at 500 nM and verapamil at 10 µM; both the stain and verapamil were added to pre-warmed imaging media and mixed vigorously to ensure full dissolution before cell application. A layer of mineral oil was applied atop the imaging media to prevent evaporation during live cell microscopy.

**DNA constructs and transfections**

DNA constructs were generated using standard cloning procedures, typically by Gibson assembly reactions and T4 ligations. Inserts were generated by PCR amplification or acquired as custom-synthesized DNA fragments (Integrated DNA Technologies, IDT). Plasmid backbones were linearized by digestion with restriction enzymes. Cloning of lentiviral backbones and sequences containing repeat regions was performed using NEB Stable Competent *e. coli* cells as the transformation host.

DNA transfections were carried out using Lipofectamine 3000 Reagent (L3000001, ThermoFisher) according to the manufacturer's instructions. For analyses involving transient transfection of HEK293-FT derived reporter cells (UAS:H2B-mCherry), 10 ng of receptor-encoding plasmid DNA was used in combination with 10 ng of a separate plasmid encoding a mTurquoise2 fluorescent protein as a co-transfection marker, combined with 30 ng of salmon sperm filler DNA. Mixtures containing 50 ng of total DNA were used to transfect ~60,000 reporter cells to be plated in a 96-well. In cases where receptor-T2A-BFP fusions were used, ~60,000 reporter cells were transfected with 10 ng of receptor-T2A-BFP plasmid with 40 ng of salmon sperm filler DNA.

**HCR-FISH**

An oligonucleotide probe set against the *mCherry* transgene mRNA was obtained through Molecular Instruments as part of an "HCR v3.0" kit. Detection was performed using "B1" amplifiers as AlexaFluor647 conjugates. Cells were fixed at room temperature with a 4% paraformaldehyde solution in Dulbecco's PBS (DPBS) for 10 minutes, were washed twice with DPBS for 5 minutes each, and permeabilized in ice-cold 70% ethanol (v/v in water) with incubation overnight at -20° C. The manufacturer's protocol was followed for probe hybridization and hybridization chain reaction (HCR) steps.

**KO cell line generation**

Cas9-mediated KO of PSEN1 and PSEN2 were done using the sgRNA provided below. Annealed oligonucleotides corresponding to the sequences were cloned into the pGuide-it-tdTomato vector backbone (Takara, 632604) according to the manufacturer's protocol.

hPSEN1-sgRNA: 5'-CCCTGTGACTCTCTGCATGG-3'
hPSEN2-sgRNA: 5'-GAAGAGCTGACCCTCAAATA-3'

HEK293-FT reporter cells (UAS:H2B-mCherry) were transfected in a 6-well by Lipofectamine using the manufacturer's protocol (ThermoFisher). At 24 hours post-transfection, tdTomato+ cells were sorted into a 96-well plate at one cell per well using a FACSMelody Cell Sorter (BD Biosciences). PSEN2 KO cells were used to generate the DKO line.

**Lentiviral production and transduction**

For viral transduction, lentiviral particles were generated using a second-generation lentiviral vector system. HEK293-FT cells were grown to approximately 90% confluence on a fibronectin-coated 6-well dish (coated using a 1 µg/mL fibronectin coating solution in PBS). The wells were transfected with 566 ng of construct-encoding transfer plasmid, alongside 833 ng of packaging (pPax2) and envelope encoding (pVSVG) plasmids using Lipofectamine 3000 Reagent.

Transfection media was replaced the following morning with 2 mL of fresh media. The viral supernatant was harvested 24 hours later and passed through a low protein-binding 0.45 μm filter before immediate use or storage at -80C. For viral transduction, cells were transduced in growth media containing diluted and filtered viral supernatant; viral transduction media was replaced with fresh media 24 h later.

**Reporter cell line generation**

U2OS:TRE3G-mCherry, and C3H/10T½:TRE-DsRed reporter cells were generated in a similar manner as was described previously (Sloas et al. 2023). Briefly, lentiviral particles were generated as described above and were used to transduce ~75,000 of U2OS or C3H/10T½ cells in a 24-well plate. Viral media was replaced with fresh media 24 hours after transduction. At 48 – 72 hours post-transduction, cells were trypsinized and transferred to a 6-well for selection in antibiotic-containing media (0.5 μg/mL puromycin for the utilized reporter constructs). Care was taken to prevent C3H/10T½ from reaching confluence. Selection in puromycin typically proceeded for 10-14 days, after which U2OS:TRE3G-mCherry cells were used a stable pool. A single clone of C3H/10T½:TRE-DsRed2 cells via limited dilution into 96 well plates before use.  The activities of the U2OS pool and of the isolated C3H/10T½ clone were verified by overnight doxycycline treatment (~200-1000 ng/uL) followed by fluorescence microscopy detection of fluorescent protein expression. Cell lines bearing doxycycline-inducible reporter genes were maintained in media containing doxycycline/tetracycline-free FBS.

**Receptor-expressing cell line generation**

Stable receptor-expressing cell lines were generated by lentiviral integration of EF1A-driven constructs. Viral particles generated as described above were used to transduce HEK293-FT:UAS-H2B-mCherry (clone E5) and U2OS:UAS-T2A-DsRed2 (clone 1G4) reporter cells developed in (Sloas et al. 2023). Typically, ~300,000 HEK293-FT and ~150,000 U2OS cells were transduced with 500 μL of viral supernatant in a 6-well. Viral media was replaced with fresh media 24 hours later. Once the cells reached ~75% confluence, media was replaced with antibiotic selection media (75 μg/mL and 200 μg/mL hygromycin-B-gold for HEK293-FT and U2OS, respectively). Cells were sub-passaged once 90% confluent, using 0.25% trypsin solution without EDTA. Media was replaced with fresh antibiotic-containing media daily, or every other day to generate receptor-expressing stable pools; selection typically proceeded for 10 – 14 days, with trypsinization and dilution as needed. Clonal cell lines were generated by limiting dilution in 96-well plates. Important note: use of 0.25% trypsin solution without EDTA is required to maintain receptor quiescence during trypsinization and sub-passaging (EDTA induces NRR unfolding and receptor cleavage; Rand et al. 2000).

For generating $\alpha$FITC(E2)-SynNotch-TetR-VP48 cells, a lentiviral construct encoding an $\alpha$FITC-SynNotch receptor with a TetR-VP48 ICD (a.k.a tTA) (pHR-SFFV-$\alpha$FITC(E2)-SynNotch-TetR-VP48) was produced in HEK293-FT cells grown in tetracycline (Tet)-free media. Transductions were performed in fibronectin (FN)-coated 24-well tissue culture plates using 100k C3H/10T½ or U2OS TRE reporter cells per well in combination with 200 µL of viral supernatant from frozen stocks stored at -80° C. After 24 hours, the cells were trypsinized using a pre-warmed 0.25% trypsin solution (without EDTA) and transferred to 6 well tissue culture plates before further expansion or analysis.

For generating $\alpha$FITC(E2)-SynNotch-Gal4-VP64 and $\alpha$FITC(E2)-SynNotch-Gal4-VP64-mTurq2 cells, HEK293-FT (UAS:H2B-mCherry) cells were transfected with linearized versions of corresponding receptor plasmids based on restriction enzyme cleaved pcDNA3 plasmids. The plasmids contained a hygromycin resistance expressed from a PGK promoter. The PGK promoter was used in place of an original SV40 promoter/origin within the pcDNA3 backbone due to the expression of the Large T-antigen in HEK293-FT cells. Plasmids were linearized by enzyme digestion at positions outside of the receptor and resistance marker cassettes, and the resulting linearized DNA was transfected into HEK293-FT (UAS:H2B-mCherry) cells. At 72 h post-transfection, stable integrants were selected using 75 µg/mL Hygromycin-B-Gold (InvivoGen). Single clones were isolated by limited dilution into 96-well plates. The clonal lines were maintained in media containing 100 µg/mL zeocin (for maintenance of the UAS:H2B-mCherry reporter) and 75 µg/mL Hygromycin-B-Gold (for maintenance of the integrated receptor encoding gene cassettes).

**Measurement of receptor surface expression and analysis of furin/S1-processing**

For cell surface measurements by flow cytometry, ~60,000 HEK293-FT cells were transfected using Lipofectamine 3000 as the transfection reagent; 100 ng of a receptor-encoding plasmid and 10 ng of pcDNA3-mTurquoise2 as a co-transfection marker. Cells were immunostained at ~24 hours after transfection using mouse monoclonal anti-c-myc-AlexaFluor647 diluted into growth medium (1:200 dilution). Staining proceeded for 30 min at 37° C before removing antibody-containing mediate and washing gently with fresh media (3 x 200 µL washes).

Cells were seeded in FN-coated 8-well glass-bottom imaging dishes to detect cell surface-expressed receptors by fluorescence imaging. HEK293-FT and CHO-K1 cells were seeded at 120k cells per well and U2OS cells at 75k cell per well. Cells were co-transfected 5-6 hours later using a mixture containing receptor-encoding plasmid and pcDNA3-mTurquoise2 co-transfection marker (50 ng of each plasmid per well). A no-receptor control was carried out by transfecting cells with 50 ng of pcDNA3-mTurquoise2 in combination with 50 ng of salmon sperm DNA as filler. Cells were stained 24 hours after transfection using anti-myc-AlexaFluor647 as described above.

HEK293-FT and U2OS cells were transfected for western blotting using conditions described for live cell imaging. Lysates were prepared by rinsing cells once with PBS, followed by direct lysis in 1xSDS-PAGE sample buffer. Samples were subjected to sonication to shear genomic DNA and reduce sample viscosity. Samples were heated and reduced before separation on SDS-PAGE gels. Proteins were transferred to nitrocellulose membranes and subsequently blocked in a “blocking buffer” solution containing 5% nonfat dry milk (w/v) in PBS containing 0.1% (v/v) Tween-20 (PBS-T). Myc-tagged receptors were detected using rabbit monoclonal anti-myc (Cell Signaling Technology), probing overnight at 4° C at a dilution of 1:1000 in blocking buffer. Anti-rabbit-HRP was used as a secondary antibody, probing at room temperature for 1 h with agitation at a dilution of 1:3000 in PBS-T. Membranes were developed using SuperSignal West Pico PLUS Chemiluminescent Substrate with recording on a ThermoFisher iBright instrument. Developed membranes were stripped using Restore Western Blot Stripping Buffer (ThermoFisher) per the supplier’s protocol, and stripped membranes were re-blocked with blocking buffer for 1 h before re-probing. The detection of loading control proteins (using anti-GAPDH, etc.) was done after detecting the primary antigens of interest. Loading control proteins were detected using stripped and re-blocked membranes with probing via DirectBlot anti-GAPDH-HRP (BioLegend). Gels were run with PageRuler Prestained Protein Ladder (10 to 180 kDa, ThermoFisher) as the molecular weight marker.

**Plating fluorescein ligands**

Ligand-coated surfaces for BSA conjugated fluorescein derivatives (BSA-5-FITC, BSA-5-fluorescein, BSA-6-fluorescein) were prepared by diluting the BSA ligand into PBS containing 10 µg/mL fibronectin. Non-TC treated wells (50 µL for 96-well, 100 µL for 8-well) were incubated for 1 hour and washed 3 times with PBS before seeding cells; wells were coated using BSA conjugate solutions at 5 µg/mL, unless otherwise indicated. Gelatin ligand-coated surfaces (gelatin-5-FITC, gelatin-5-OG488) were plated similarly using 5 µg/mL solutions in combination with 50 µg/mL unconjugated gelatin in PBS.

**Generation of BSA-5-fluorescein and BSA-6-fluorescein ligands**

A 10:1 molar ratio of NHS-ester-containing molecule to BSA was used and the reaction was carried out for 2 hrs at room temperature in 100 mM Borate Buffer (pH 8.4). Each reaction mixture was buffer exchanged into PBS using a 2 KDa MWCO Slide-A-Lyzer Dialysis Cassette (ThermoFisher). Buffer exchanged BSA-DBCO was then reacted with excess amounts of 6-FAM-azide and 5-FAM azide.

**Transactivation coculture assay**

For coculture experiments, ~20,000 HEK293-FT-based “sender” cells expressing anti-bio-SNAP-TMD-DLL1 ligand were combined with ~20,000 HEK293-FT (UAS:H2B-mCherry)-based “receiver” cells expressing myc-LaG16-$\alpha$FITC(E2)-SynNotch-Gal4-VP64-T2A-BFP. Suspended cell mixtures were then added to fibronectin-coated wells in a 96-well plate. At ~6 hours post-cell seeding (once cells had adhered), the cocultures were treated with bifunctional bridging compounds at the concentrations indicated within the captions. Cocultures containing non-transduced HEK293-FT cells in place of ligand-expressing senders were used as controls; control cocultures were seeded into microwells and treated with bridging compounds in a similar manner as described for sender-receiver mixtures.

**Preparation and cell stimulation with ligand-decorated beads**

Fluorescein-bound microparticles were prepared using streptavidin-coated magnetic beads (Dynabeads M-280 Streptavidin, 2.8-µm, ThermoFisher 60210). Beads were first rinsed by diluting 10 µL of the commercial suspension into 1 mL PBS. The beads were then collected by magnetic separation and resuspended in 1 mL PBS solution containing 200 nM biotin-FITC or biotin-OG488. The mixture was allowed to incubate for 30 min at room temperature, after which they were magnetically collected and rinsed three times using PBS to remove unbound ligand molecules. A fourth rinse was carried out using 2 mL of cell culture media. The beads were resuspended in a 130 µL volume of culture media before cell application. Based on this protocol, the final mixture corresponded to ~500k beads per 10 µl of fully suspended solution; beads were applied to cells using 10 µl volumes per each well of an 8-well imaging dish (~500k beads per well).

**Tetrazine-TCO ligation-mediated gene expression control**

TCO-BSA protein conjugation was prepared as described by (McMahan and Ngo 2022) using TCO-PEG4-NHS (Click Chemistry Tools); a step-by-step protocol is provided below. Reactions were carried out using solutions containing 75 µM BSA dissolved in 75 mM sodium bicarbonate buffer (pH 8.2). Protein conjugation was initiated by adding NHS ester from stock solutions dissolved in dry DMSO. Final reaction solutions contained 7.5 mM of the NHS ester and 20% (v/v) DMSO. Coupling reactions were carried out for 1 h at room temperature, followed by removal of unconjugated and hydrolyzed esters via three rounds of dialysis against PBS using 10 kDa MWCO filters (ThermoFisher). Following dialysis, the solution was sterilized by filtration through a 0.2 µm porous membrane filter.

TCO-BSA coated surfaces were prepared in a similar manner as described above for BSA-fluorescein ligands using 5 µg/mL coating solutions. Receptor-expressing cells were seeded on

TCO-BSA- coated substrates and allowed to adhere at 37° C for ~6 hours. Cells were exchanged into fresh media before treatment with Tz-5-fluorescein at the concentrations indicated within the captions. Reporter expression analyses were conducted at 24 hours post-treatment.

For live timelapse microscopy of Tz-TCO-mediated gene expression, cells were grown in coverslip bottom imaging dishes for recordings via an inverted epifluorescence microscope. Neutravidin was used to immobilize a TCO-$PEG_4$-biotin reagent as follows: wells were pre-coated with fibronectin and biotinylated-BSA using a solution containing 20 μg/mL FN and 25 μg/ml biotinylated-BSA in PBS. Coating with the solution proceeded for 1 hour at 37° C before removing the solution and rinsing three times with PBS. NeutrAvidin (NA) was then immobilized on the coated surfaces via treatment with a 100 μg/mL NA solution in PBS. NA binding proceeded for 30 minutes at room temperature before removal and rinsing three times with PBS. TCO-$PEG_4$-biotin was then immobilized on the NA-containing surfaces by treatment with a solution containing 200 nM TCO-$PEG_4$-biotin in PBS for 20 minutes at room temperature before removal and rinsing three times with PBS. Stable clones of receptor-expressing HEK293-FT(UAS:H2B-mCherry) and U2OS(UAS:dsRed2) were added to the wells and allowed to adhere for ~4 hours. Following cell attachment, Tz-5-fluorescein was diluted into the cell and media-containing wells using 2x concentrations solutions to achieve the final concentrations indicated in the captions. Negative (TCO-lacking) control wells were prepared as described above but omitting TCO-$PEG_4$-biotin. Positive control wells were prepared by treatment of NA-bound wells with 200 nM biotin-FITC (in place of TCO-$PEG_4$-biotin and tetrazine-5-FAM).

**Step-by-step protocol for preparing the light-conditional BSA-PC-5-Fluorescein ligand conjugate.**

All steps involving PC-5-fluorescein were carried out with minimized exposure to light. Preparation of the BSA-PC-5-Fluorescein conjugate was done according to the step-by-step protocol below.

1. Dissolve 3 mg of BSA in 2 mL of 1X borate buffer (pH 8.5).
2. Remove a 1 mg tube of CMNB-Caged Carboxyfluorescein, SE (succinimidyl ester) from storage at -20 ° C and briefly centrifuge to collect the tube contents at the bottom of the tube. Allow the tube to equilibrate to room temperature (~15 min).
3. Add 100 μL of dry DMSO to the tube containing CMNB-Caged Carboxyfluorescein.
4. Vortex the tube to ensure dissolution of the dye and briefly centrifuge the tube to collect its contents at the bottom.
5. Combine the 2 mL BSA solution with the 100 μL of dissolved CMNB-Caged Carboxyfluorescein and mix thoroughly.
6. Allow the reaction to proceed at room temperature for 4 hours, with protection from light.

7. At the end of the reaction, transfer the 2.1 mL reaction mixture to a Slide-A-Lyzer G2 Dialysis Cassette (3 mL capacity, 10K MWCO). Add 900 µL of borate buffer solution to bring the volume to 3 mL before capping the cassette.
8. Submerge the cassette in 2 L of PBS. Allow the mixture to equilibrate for 2 hours at room temperature, with gentle stirring and protection from light.
9. Perform a second round of dialysis as in step 8, using a fresh 2 L of PBS.
10. Perform a final round of dialysis against a fresh 2 L PBS overnight at 4 C.
11. Following dialysis, store the product as 150 µL aliquots in dark tubes. Keep at -20 ° C, with protection from light until use.

**Step-by-step protocol for coating imaging dishes with BSA-PC-5-fluorescein.**

All steps involving PC-5-Fluorescein were carried out with minimized exposure to light. Coating of BSA-PC-5-Fluorescein conjugate to imaging wells was carried out according to the step-by-step protocol below.

1. Thaw an aliquot of the 1 mg/ml BSA-PC-5-Fluorescein and prepare a 25 µg/mL working solution by diluting the conjugate into PBS containing 10 µg/mL fibronectin.
2. Coat a coverslip bottom by applying the solution to glass-bottom imaging dishes. Allow the protein conjugate to adhere to the dish's surface for 1.5 hours at 37° C, with protection from light. Note: avoid direct contact with the glass surface during all subsequent steps to avoid scratching or dislodging BSA-PC-5-Fluorescein from the coverslip bottom.
3. After the coating step, remove the solution by inversion of the dish.
4. Rinse the coat wells 3X using PBS to remove unbound protein conjugate.
5. Following the rinse steps, add enough PBS to the wells to cover the bottom surface.
6. Apply the photomask to the outer bottom of the well by dropping a small amount of mineral oil on the printed transparency. Apply the oil to the back side of the transparency (opposite of the laser-printed surface).
7. Place the dish/vessel on the oil-containing side of the transparency. Push down gently. Be sure the photomask region is positioned in the center of each well region.
8. Wick excess oil away carefully using a Kimwipe or lens paper.
9. Place the microwell atop the UV lamp. Illuminate the sample for 3 minutes.
10. Remove transparency. Carefully clean the bottom of the dish to remove the remaining oil.
11. Remove PBS. Rinse once with cell media.
12. Add 100k U2OS cells (in pre-warmed media)

**Preparation, cell seeding, and analysis of cells grown on patterned ligand surfaces**

A printed photomask was attached to the bottom of the imaging dish. A small volume of mineral oil was applied to the outer bottom of the imaging dishes before the printed photomask was applied. BSA-PC-5-Fluorescein was then photo-uncaged for 2 minutes by direct placement atop a handheld UV lamp (365 nm, 6 W, Research Products International, item # 950006-02). The wells were rinsed two times with PBS following the uncaging step. Cells were applied to the wells in media containing 10 µM DAPT. For 8 well imaging dishes, 100,000 U2OS cells were applied per well. DAPT was removed the next day by gentle aspiration of DAPT-containing media followed by gentle rinsing thrice with DAPT-free media. Cells were returned to the incubator in fresh media and imagined the next day.  Imaging to detect co-expressed T2A-BFP was done under live cell conditions. For nuclear counterstaining, imaging was performed in media containing Hoechst-JaneliaFluor646 (Hoechst-JF646) at 500 nM and containing verapamil at 10 µM; both the stain and verapamil were added to pre-warmed imaging media and mixed vigorously to ensure full dissolution before cell application. After 15-30 minutes of staining, cells were fixed with 4% paraformaldehyde (PFA, w/v, diluted into PBS from a 16% stock). Fixation proceeded for 15 minutes at 37° C, after which the PFA was removed. Cells were rinsed once with culture media (to quench trace PFA) and again with PBS.

Example of photomask printed on clear transparency film for 8-well chambered dish (well dimensions are units are in mm):

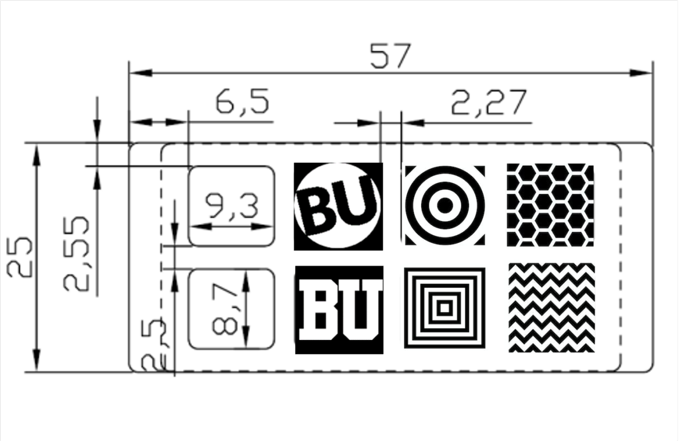


**Expression, purification, and dye-conjugation of HaloTag-CNA35**

Starter cultures of BL21-DE3 *E. coli* cells harboring pET28a Halotag-CNA35 (AddGene # 131128) were grown in 2xYT media containing 35 µg/ml kanamycin. Cultures were grown overnight with incubation at 37° C and shaking (2 x 5 mL cultures). The next day, ~10 mL of the overnight cultures were used to inoculate fresh 500 mL of 2xYT containing 35 µg/ml kanamycin. Cells were grown at 37° C with shaking (250 rpm) until a culture density of $OD_{600}$ = ~0.5 was reached, at which point the incubation temperature was adjusted to 27° C. Protein expression was induced after 15 minutes of growth at 27° C by the addition of IPTG to the culture medium to a final concentration of 0.5 mM. Expression proceeded at 27° C, shaking (250 rpm) for ~8 hours. Cells were harvested by centrifugation in 50 mL tubes. Cell pellets were stored at -80° C until use. For purification, 2 x 50 mL aliquots were processed using one column from the Ni-NTA

Fast Start Kit (Qiagen, 30600). The purity of the protein fractions was assessed by Coomassie staining (GelCode Blue Stain Reagent, Pierce), and the purest fractions were utilized in conjugation reactions with a HaloTag-reactive OG488.

A chloroalkane (CA)-OG488-biotin was prepared by reacting Halo-DBCO (Iris Biotech, RL-3670.0100) and OG488-biotin-azide as follows: a 20 mM stock solution of Halo-DBCO was prepared by dissolving 2.6 mg of dry material in 254 µL of DMSO. An aliquot of this solution was diluted to 2 mM in DMSO, and the remainder of the 20 mM stock was stored at -80° C with the labeling of the date of preparation. The 2 mM Halo-DBCO solution was then reacted with a 2 mM solution of OG488-biotin-azide, also in DMSO. The reaction was initiated by combining 40 µL of Halo-DBCO with 50 µL of OG488-biotin-azide; the excess OG488-biotin-azide was used to ensure complete derivatization of Halo-DBCO. The solution was incubated at room temperature for 3 hours before use. For protein conjugation, the reaction mixture was diluted 1:250 by volume into aliquots of purified HaloTag-CNA35 in the Ni-NTA Fast Start Kit elution buffer. The solution was mixed by pipetting, and the reaction proceeded at 4° C for three hours before unligated components were removed from the solution by filtration through 10 kDa MWCO filters (66380, ThermoFisher); the proteins were then exchanged into 50 mM Tris buffer (pH = 8.0) by repeated centrifugation and dilution of the retentate. The conjugate concentration was determined via extinction coefficient (OG488, $\epsilon$ = 76,000 $cm^{-1}M^{-1}$ at 495 nm). Solutions of the conjugated protein (containing 12 µM of product) were then frozen by immersion in liquid nitrogen followed by storage at -20° C until use. The purity of the isolated protein and confirmation of its successful dye-conjugation were confirmed via SDS-PAGE. Lanes containing dye-conjugated and unconjugated (control) protein samples were compared by in-gel fluorescence detection. The gel was fixed in a fixing solution (50% methanol, 40% water, 10% glacial acetic acid) and rinsed with water before total protein gel fluorescence was detected using an iBright instrument (ThermoFisher). The gel was then stained for total protein using the Coomassie-based GelCode Blue Stain Reagent (Pierce).

**Collagen adsorption, OG488-CNA-binding, and cell stimulation**

A non-tissue culture (non-TC)-treated 96-well plate (Celltreat, 229596) was coated with native collagen-I using a 50 µg/mL solution diluted from a 4.04 mg/mL solution provided by the supplier. Coating proceeded for 1 hour at 20 C, after which the collagen solution was gently removed, and wells were rinsed 3 times using PBS. The coated wells were then passivated using a 1% solution of BSA (w/v) in PBS; passivation proceeded for 1 hour at room temperature before gently aspirating the BSA solution and rinsing three times PBS. The collagen-coated and BSA-passivated wells were then treated with varying concentrations of CNA-OG488 (as indicated in the figures and captions); CNA binding proceeded for 1 hour at room temperature before gentle aspiration of the CNA-containing solution and rinsing 3 times with PBS. Cells suspended in pre-warmed (37° C) growth media were then added to the treated wells at 60k cells per well. Flow cytometry measurement of reporter activities was carried out 24 hours after cell plating. Control wells coated with either fibronectin (FN) or gelatin were prepared using coating solutions of 10 µg/mL in PBS (for FN) and 50 µg/mL gelatin in PBS. FN and gelatin-coated wells were also treated with OG488-CNA as described above, and cells grown in these wells were analyzed as CNA non-binding controls.

**Collagen denaturation, CHP-fluorescein binding, and cell stimulation**

A non-tissue culture (non-TC)-treated 96-well plate (Celltreat, 229596) was coated with a 50 µg/mL solution of native collagen-I followed by washing with PBS; a protocol similar to that which is described above was used. To produce wells containing denatured collagen, microwells containing adhered native collagen were subjected to heat denaturation by rinsing wells with sterile water ($ddH_2O$) pre-headed to ~90° C. Three rinses with boiling hot $ddH_2O$ were carried out to denature plated collagen immediately before treatment with CHP-fluorescein. To produce control wells containing native collagen, wells were treated in a similar manner but rinsed using room-temperature sterile $ddH_2O$ (in place of hot $ddH_2O$). The denatured and control wells were then treated with CHP-fluorescein. A stock solution containing 50 µM CHP-fluorescein in PBS was prepared according to the manufacturer's protocol; aliquots from this stock were further diluted to PBS to generate binding solutions at the working concentrations indicated in the figures and captions. For CHP-fluorescein binding, peptide working solutions were applied to native and denatured collagen-containing wells at a volume of 50 µL per well. Wells were incubated with the solutions for 3 hours at room temperature before gently removing the peptide-containing solutions and washing wells 3 times using room temperature PBS. Cells suspended in pre-warmed growth media were then added to the treated wells at 60k cells per well. Flow cytometry measurement of reporter levels was carried out 24 hours after cell plating.

**Myogenic differentiation of C3H/10T½ cells.**

A clonal cell line containing an integrated TRE:p65-MyoD-T2A-DsRed construct was utilized. Generation and isolation of the clone have been previously described (Sloas et al. 2023). The DNA construct used in generating the line has also been described (Kabadi et al., 2014; AddGene #60627). A control line containing an inducible TRE:DsRed2 reporter (without p65-MyoD) was also generated (Kabadi et al., 2014; AddGene #60623). For experiments using C3H/10T½ reporter lines, cells were transduced with lentiviral constructs based on pHR-SFFV-$\alpha$FITC(E2)-SynNotch-TetR-VP48.

Experiments using transduced C3H/10T½ cells were typically initiated at 48-72 hours post-transduction as follows: for surface detection of $\alpha$FITC(E2)-SynNotch expression in transduced C3H/10T½ cells, cells were plated in FN-coated 8 well coverslip bottom imaging dishes. Antibody staining under live cell conditions was carried out the next day using mouse anti-myc-AlexaFluor647 antibody diluted into tetracycline-free culture media (1:200). Staining was carried for 30 minutes at 37° C. Cells were washed with pre-warmed culture media (3 x 200 µL washes) and counterstained with Hoechst 33342 in media for 30 minutes at 37° C. Imaging was done under live cell conditions in PBS. Non-transduced C3H/10T½ cells were stained and imaged as a labeling control.

For bio-orthogonal Tz/TCO-ligation-induced differentiation, 18 well glass bottom imaging dishes were treated with a PBS solution containing 20 µg/ml FN and 50 µg/mL TCO-BSA for 3 hours at 37° C. Wells were rinsed three times with PBS before adding cells to a density of 30k C3H/10T½ cells per well. After 2 hours, wells were treated with Tz-5-fluorescein by adding dye-containing media solution at 2X concentrations to achieve the final Tz-5-fluorescein indicated in the figures and captions.

For patterned differentiation, 8 well glass-bottom imaging dishes were coated with BSA-PC-5-Fluorescein and FN, as described above. Photo-uncaging was carried out for 2 minutes, and wells were rinsed with PBS before adding cells. Transduced C3H/10T½ cells were trypsinized using pre-warmed 0.25% trypsin solution (without EDTA) and quenched with media before collection by centrifugation (5 minutes at 300 x g). Cells were resuspended in fresh media before seeding into the photo-patterned 8-well dishes. For C3H/10T½, ~65,000 cells were seeded per well of patterned 8-well dishes. Cells were analyzed at 2 - 3 days after seeding. See below regarding the immunofluorescence detection of myosin heavy chain expression in differentiated cells.

To verify myosin heavy chain (MHC) expression, cells were fixed using 4% paraformaldehyde (PFA, w/v, diluted into PBS from a 16% stock). Fixation proceeded for 15 minutes at 37° C, after which the PFA was removed. Cells were rinsed once with culture media (to quench trace PFA) and again with PBS. Cells were permeabilized with a PBS solution containing 0.1% Triton-X100 (v/v) (10 minutes at room temperature), rinsed with PBS, and subsequently blocked using Immunofluorescence Blocking Buffer (Cell Signaling Technologies) for 60 minutes at room temperature. Antibody staining was done with mouse anti-myosin heavy chain antibody at a 1:100 dilution in antibody dilution buffer (PBS containing 0.1% Tween-20, v/v, and 1% BSA, w/v; sterile filtered before use). Staining with the primary antibody proceeded overnight at 4° C. The primary antibody solution was removed the next day and washed using PBS-T thrice for 10 minutes each. Cells were then labeled with a rabbit anti-mouse AlexaFluor647-conjugate using a 1:1000 dilution in antibody dilution buffer for 1 hour at room temperature. The unbound secondary antibody was removed by washing with PBS-T three times for ten minutes each. Cells were counterstained with Hoechst 33342 before imaging in PBS.

For CHP-5-fluorescein myogenic conversion of C3H/10T½ fibroblast cells, 8 well glass-bottom imaging dishes were coated with 50 µg/ml collagen rinsed thrice with PBS, and heat-denatured as described above. A solution containing 2 µM CHP-5-fluorescein was applied to the collagen-containing wells, and unbound units were removed by rinsing (both as described above). C3H/10T½ TRE:p65-MyoD-2A-DsRed2 cells were seeded into the treated wells at ~65,000 cells per well.

**Protein Sequences:**

**SS-myc-anti-FITC(E2)-SynNotch-Gal4VP64-mTurquoise2**

MALPVTALLLPLALLLHAARPEQKLISEEDLAAQVQLVESGGNLVQPGGSLRLSCAASGFTFGS
FSMSWVRQAPGGGLEWVAGLSARSSLTHYADSVKGRFTISRDNAKNSVYLQMNSLRVEDTAV
YYCARRSYDSSGYAGHFYSYMDVWGQGTLVTVSGGGGSGGGGSGGGGSSVLTQPSSVSAA
PGQKVTISCSGSTSNIGNNYVSWYQQHPGKAPKLMIYDVSKRPSGVPDRFSGSKSGNSASLDI
SGLQSEDEADYYCAAWDDSLSEFLFGTGTKLTVLGILDYSFTGGAGRDIPPPQIEEACELPECQ
VDAGNKVCNLQCNNHACGWDGGDCSLNFNDPWKNCTQSLQCWKYFSDGHCDSQCNSAGC
LFDGFDCQLTEGQCNPLYDQYCKDHFSDGHCDQGCNSAECEWDGLDCAEHVPERLAAGTLV
LVVLLPPDQLRNNSFHFLRELSHVLHTNVVFKRDAQGQQMIFPYYGHEEELRKHPIKRSTVGW
ATSSLLPGTSGGRQRRELDPMDIRGSIVYLEIDNRQCVQSSSQCFQSATDVAAFLGALASLGSL
NIPYKIEAVKSEPVEPPLPSQLHLMYVAAAAFVLLFFVGCGVLLSRKRRRMKLLSSIEQACDICR
LKKLKCSKEKPKCAKCLKNNWECRYSPKTKRSPLTRAHLTEVESRLERLEQLFLLIFPREDLDMI
LKMDSLQDIKALLTGLFVQDNVNKDAVTDRLASVETDMPLTLRQHRISATSSSEESSNKGQRQL
TVSAAAGGSGGSGGSDALDDFDLDMLGSDALDDFDLDMLGSDALDDFDLDMLGSDALDDFDL
DMLGSGGGGSMVSKGEELFTGVVPILVELDGDVNGHKFSVSGEGEGDATYGKLTLKFICTTGK
LPVPWPTLVTTLSWGVQCFARYPDHMKQHDFFKSAMPEGYVQERTIFFKDDGNYKTRAEVKF
EGDTLVNRIELKGIDFKEDGNILGHKLEYNYFSDNVYITADKQKNGIKANFKIRHNIEDGGVQLAD
HYQQNTPIGDGPVLLPDNHYLSTQSKLSKDPNEKRDHMVLLEFVTAAGITLGMDELYK*

**SS-myc-anti-FITC(E2)-SynNotch-TetR-VP48**

MALPVTALLLPLALLLHAARPEQKLISEEDLAAQVQLVESGGNLVQPGGSLRLSCAASGFTFGS
FSMSWVRQAPGGGLEWVAGLSARSSLTHYADSVKGRFTISRDNAKNSVYLQMNSLRVEDTAV
YYCARRSYDSSGYAGHFYSYMDVWGQGTLVTVSGGGGSGGGGSGGGGSSVLTQPSSVSAA
PGQKVTISCSGSTSNIGNNYVSWYQQHPGKAPKLMIYDVSKRPSGVPDRFSGSKSGNSASLDI
SGLQSEDEADYYCAAWDDSLSEFLFGTGTKLTVLGILDYSFTGGAGRDIPPPQIEEACELPECQ
VDAGNKVCNLQCNNHACGWDGGDCSLNFNDPWKNCTQSLQCWKYFSDGHCDSQCNSAGC
LFDGFDCQLTEGQCNPLYDQYCKDHFSDGHCDQGCNSAECEWDGLDCAEHVPERLAAGTLV
LVVLLPPDQLRNNSFHFLRELSHVLHTNVVFKRDAQGQQMIFPYYGHEEELRKHPIKRSTVGW
ATSSLLPGTSGGRQRRELDPMDIRGSIVYLEIDNRQCVQSSSQCFQSATDVAAFLGALASLGSL
NIPYKIEAVKSEPVEPPLPSQLHLMYVAAAAFVLLFFVGCGVLLSRKRRRQLCIQKLMSRLDKSK
VINSALELLNEVGIEGLTTRKLAQKLGVEQPTLYWHVKNKRALLDALAIEMLDRHHTHFCPLEGE
SWQDFLRNNAKSFRCALLSHRDGAKVHLGTRPTEKQYETLENQLAFLCQQGFSLENALYALSA
VGHFTLGCVLEDQEHQVAKEERETPTTDSMPPLLRQAIELFDHQGAEPAFLFGLELIICGLEKQL
KCESGGPADALDDFDLDMLPADALDDFDLDMLPADALDDFDLDMLPG*

**SS-myc-anti-FITC(E2)-SynNotch-Gal4VP64**

MALPVTALLLPLALLLHAARPEQKLISEEDLAAQVQLVESGGNLVQPGGSLRLSCAASGFTFGS
FSMSWVRQAPGGGLEWVAGLSARSSLTHYADSVKGRFTISRDNAKNSVYLQMNSLRVEDTAV
YYCARRSYDSSGYAGHFYSYMDVWGQGTLVTVSGGGGSGGGGSGGGGSSVLTQPSSVSAA
PGQKVTISCSGSTSNIGNNYVSWYQQHPGKAPKLMIYDVSKRPSGVPDRFSGSKSGNSASLDI
SGLQSEDEADYYCAAWDDSLSEFLFGTGTKLTVLGILDYSFTGGAGRDIPPPQIEEACELPECQ
VDAGNKVCNLQCNNHACGWDGGDCSLNFNDPWKNCTQSLQCWKYFSDGHCDSQCNSAGC
LFDGFDCQLTEGQCNPLYDQYCKDHFSDGHCDQGCNSAECEWDGLDCAEHVPERLAAGTLV
LVVLLPPDQLRNNSFHFLRELSHVLHTNVVFKRDAQGQQMIFPYYGHEEELRKHPIKRSTVGW
ATSSLLPGTSGGRQRRELDPMDIRGSIVYLEIDNRQCVQSSSQCFQSATDVAAFLGALASLGSL
NIPYKIEAVKSEPVEPPLPSQLHLMYVAAAAFVLLFFVGCGVLLSRKRRRMKLLSSIEQACDICR
LKKLKCSKEKPKCAKCLKNNWECRYSPKTKRSPLTRAHLTEVESRLERLEQLFLLIFPREDLDMI
LKMDSLQDIKALLTGLFVQDNVNKDAVTDRLASVETDMPLTLRQHRISATSSSEESSNKGQRQL
TVSAAAGGSGGSGGSDALDDFDLDMLGSDALDDFDLDMLGSDALDDFDLDMLGSDALDDFDL
DMLGS*

**SS-myc-4M5.3-SynNotch-Gal4VP64**

MALPVTALLLPLALLLHAARPEQKLISEEDLAADVVMTQTPLSLPVSLGDQASISCRSSQSLVHS
NGNTYLRWYLQKPGQSPKVLIYKVSNRVSGVPDRFSGSGSGTDFTLKINRVEAEDLGVYFCSQ
STHVPWTFGGGTKLEIKSSADDAKKDAAKKDDAKKDDAKKDGGVKLDETGGGLVQPGGAMKL
SCVTSGFTFGHYWMNWVRQSPEKGLEWVAQFRNKPYNYETYYSDSVKGRFTISRDDSKSSV
YLQMNNLRVEDTGIYYCTGASYGMEYLGQGTSVTVSILDYSFTGGAGRDIPPPQIEEACELPEC
QVDAGNKVCNLQCNNHACGWDGGDCSLNFNDPWKNCTQSLQCWKYFSDGHCDSQCNSAG
CLFDGFDCQLTEGQCNPLYDQYCKDHFSDGHCDQGCNSAECEWDGLDCAEHVPERLAAGTL
VLVVLLPPDQLRNNSFHFLRELSHVLHTNVVFKRDAQGQQMIFPYYGHEEELRKHPIKRSTVG
WATSSLLPGTSGGRQRRELDPMDIRGSIVYLEIDNRQCVQSSSQCFQSATDVAAFLGALASLG
SLNIPYKIEAVKSEPVEPPLPSQLHLMYVAAAAFVLLFFVGCGVLLSRKRRRMKLLSSIEQACDIC
RLKKLKCSKEKPKCAKCLKNNWECRYSPKTKRSPLTRAHLTEVESRLERLEQLFLLIFPREDLD
MILKMDSLQDIKALLTGLFVQDNVNKDAVTDRLASVETDMPLTLRQHRISATSSSEESSNKGQR
QLTVSAAAGGSGGSGGSDALDDFDLDMLGSDALDDFDLDMLGSDALDDFDLDMLGSDALDDF
DLDMLGS*

**SS-myc-FluA-SynNotch-Gal4VP64**

MALPVTALLLPLALLLHAARPEQKLISEEDLDVYHDGACPEVKPVDNFDWSQYHGKWWEVAKY
PSPNGKYGKCGWIEYTPEGKSVKVSRYDVIHGKEYFMEGTAYPVGDSKIGKIYHSRTVGGYTK
KTVFNVLSTDNKNYIIGYTCRYDEDKKGHWDHVWVLSRSMVLTGEAKTAVENYLIGSPVVDSQ
KLVYSDFSEAACKVNNILDYSFTGGAGRDIPPPQIEEACELPECQVDAGNKVCNLQCNNHACG

WDGGDCSLNFNDPWKNCTQSLQCWKYFSDGHCDSQCNSAGCLFDGFDCQLTEGQCNPLYD
QYCKDHFSDGHCDQGCNSAECEWDGLDCAEHVPERLAAGTLVLVVLLPPDQLRNNSFHFLRE
LSHVLHTNVVFKRDAQGQQMIFPYYGHEEELRKHPIKRSTVGWATSSLLPGTSGGRQRRELDP
MDIRGSIVYLEIDNRQCVQSSSQCFQSATDVAAFLGALASLGSLNIPYKIEAVKSEPVEPPLPSQ
LHLMYVAAAAFVLLFFVGCGVLLSRKRRRMKLLSSIEQACDICRLKKLKCSKEKPKCAKCLKNN
WECRYSPKTKRSPLTRAHLTEVESRLERLEQLFLLIFPREDLDMILKMDSLQDIKALLTGLFVQD
NVNKDAVTDRLASVETDMPLTLRQHRISATSSSEESSNKGQRQLTVSAAAGGSGGSGGSDAL
DDFDLDMLGSDALDDFDLDMLGSDALDDFDLDMLGSDALDDFDLDMLGS*

**SS-myc-4M5.3-scFab-SynNotch-Gal4VP64**

MALPVTALLLPLALLLHAARPEQKLISEEDLAADVVMTQTPLSLPVSLGDQASISCRSSQSLVHS
NGNTYLRWYLQKPGQSPKVLIYKVSNRVSGVPDRFSGSGSGTDFTLKINRVEAEDLGVYFCSQ
STHVPWTFGGGTKLEIKRADAAPTVSIFPPSSEQLTSGGASVVCFLNNFYPKDINVKWKIDGSE
RQNGVLNSWTDQDSKDSTYSMSSTLTLTKDEYERHNSYTCEATHKTSTSPIVKSFNRNECGG
SSGSGSGSTGTSSSGTGTSAGTTGTSASTSGSGSGGGGGSGGGGSAGGTATAGASSGSGV
KLDETGGGLVQPGGAMKLSCVTSGFTFGHYWMNWVRQSPEKGLEWVAQFRNKPYNYETYY
SDSVKGRFTISRDDSKSSVYLQMNNLRVEDTGIYYCTGASYGMEYLGQGTSVTVSAAKTTPPS
VYPLAPGSAAQTNSMVTLGCLVKGYFPEPVTVTWNSGSLSSGVHTFPAVLQSDLYTLSSSVTV
PSSTWPSETVTCNVAHPASSTKVDKKIVPRILDYSFTGGAGRDIPPPQIEEACELPECQVDAGN
KVCNLQCNNHACGWDGGDCSLNFNDPWKNCTQSLQCWKYFSDGHCDSQCNSAGCLFDGF
DCQLTEGQCNPLYDQYCKDHFSDGHCDQGCNSAECEWDGLDCAEHVPERLAAGTLVLVVLL
PPDQLRNNSFHFLRELSHVLHTNVVFKRDAQGQQMIFPYYGHEEELRKHPIKRSTVGWATSSL
LPGTSGGRQRRELDPMDIRGSIVYLEIDNRQCVQSSSQCFQSATDVAAFLGALASLGSLNIPYKI
EAVKSEPVEPPLPSQLHLMYVAAAAFVLLFFVGCGVLLSRKRRRMKLLSSIEQACDICRLKKLK
CSKEKPKCAKCLKNNWECRYSPKTKRSPLTRAHLTEVESRLERLEQLFLLIFPREDLDMILKMD
SLQDIKALLTGLFVQDNVNKDAVTDRLASVETDMPLTLRQHRISATSSSEESSNKGQRQLTVSA
AAGGSGGSGGSDALDDFDLDMLGSDALDDFDLDMLGSDALDDFDLDMLGSDALDDFDLDML
GS*

**HisTag-HaloTag-CNA35**

MGSSHHHHHHSSGLVPRGSHMASMAEIGTGFPFDPHYVEVLGERMHYVDVGPRDGTPVLFL
HGNPTSSYVWRNIIPHVAPTHRCIAPDLIGMGKSDKPDLGYFFDDHVRFMDAFIEALGLEEVVL
VIHDWGSALGFHWAKRNPERVKGIAFMEFIRPIPTWDEWPEFARETFQAFRTTDVGRKLIIDQN
VFIEGTLPMGVVRPLTEVEMDHYREPFLNPVDREPLWRFPNELPIAGEPANIVALVEEYMDWLH
QSPVPKLLFWGTPGVLIPPAEAARLAKSLPNCKAVDIGPGLNLLQEDNPDLIGSEIARWLSTLEIS
GSGEFHGSARDISSTNVTDLTVSPSKIEDGGKTTVKMTFDDKNGKIQNGDMIKVAWPTSGTVKI
EGYSKTVPLTVKGEQVGQAVITPDGATITFNDKVEKLSDVSGFAEFEVQGRNLTQTNTSDDKV

ATITSGNKSTNVTVHKSEAGTSSVFYYKTGDMLPEDTTHVRWFLNINNEKSYVSKDITIKDQIQG
GQQLDLSTLNINVTGTHSNYYSGQSAITDFEKAFPGSKITVDNTKNTIDVTIPQGYGSYNSFSINY
KTKITNEQQKEFVNNSQAWYQEHGKEEVNGKSFNHTVHNINANAGIEGTVKGELKVLKQDKDT
KASVDL
